# Structural insights into human recoverin

**DOI:** 10.64898/2026.03.20.713130

**Authors:** Christopher O. MacCarthy, Alisa A. Vologzhannikova, Anatolii S. Belousov, Natalia N. Novikova, Victoria A. Rastrygina, Marina P. Shevelyova, Mikhail L. Shishkin, Natalia G. Shebardina, Mikhail B. Shevtsov, Ivan A. Kapranov, Alexey V. Mishin, Dmitrii E. Dashevskii, Yuqi Yang, Daniil A. Fedotov, Ekaterina A. Litus, Ekaterina I. Pogodina, Dmitry V. Zinchenko, Alexander L. Trigub, Alexander V. Rogachev, Sergey N. Yakunin, Philipp S. Orekhov, Sergei E. Permyakov, Valentin I. Borshchevskiy, Evgeni Yu. Zernii

**Author notes:** Corresponding authors &.

## Abstract

Recoverin is a key calcium sensor that controls the desensitization of the visual rhodopsin by GRK1. Previous studies have traditionally been conducted on bovine protein (bRec), while data on human ortholog (hRec) remain scarce. Here, we combine X-ray crystallography, X-ray absorption spectroscopy (XANES), quantum mechanical calculations, molecular dynamics, and functional assays to provide an integrated characterization of hRec. The 2Ca^2+^-bound hRec structure was solved at 1.60 Å, showing that, unlike bRec, hRec interacts with ROS membranes at physiologically relevant submicromolar Ca^2+^ levels, due to a species-specific charge distribution that might influence membrane interactions. Both recoverins form a set of Ca^2+^/Zn^2+^-bound conformers with improved functional performance. X-ray crystallography (1.85 Å) and XANES revealed a specific tetrahedral Zn^2+^ site in 1Ca^2+^-bound hRec, the first such site reported in the NCS family. In 1Ca^2+^-bound hRec, zinc promotes the formation of active state, whereas in 2Ca^2+^-state of bRec, it significantly enhances GRK1 binding, as the latter can complement the Zn^2+^ coordination. These data refine our understanding of recoverin function in humans and highlight its role as a key link between calcium and zinc signaling in mammalian photoreceptors under normal and pathological conditions.

## 1. Introduction

Recoverin is an EF-hand calcium-binding protein expressed predominantly in vertebrate rod and cone photoreceptors. Most of recoverin is localized in the inner segments and synaptic terminals, while its small but functionally significant part undergoes light-dependent translocation to membrane discs in the outer segments, where it plays a key role in Ca^2+^-dependent regulation of visual transduction and light adaptation (for review, see ^1–3^). Such activity is based on controlling GRK1 (rhodopsin kinase), an enzyme responsible for phosphorylation and thereby desensitization of visual receptor rhodopsin ^4,5^. Mechanistically, recoverin prolongs the lifetime of the photoexcited rhodopsin by inhibiting the phosphorylation until intracellular calcium concentration declines in response to the operation of the visual cascade. As a result, it decelerates the decay of phototransduction by making it Ca^2+^-dependent and accelerates the inactivation of photoexcited rhodopsin in background light by releasing GRK1 at low intracellular Ca^2+^, which is confirmed by electrophysiological studies ^6,7^. Such effects are crucial for the normal functioning of the visual system, as they increase its sensitivity in dim light and adjust it when light levels increase, thereby preventing premature saturation.

Being a typical member of the neuronal calcium sensor (NCS) family ^8^, recoverin (23 kDa) consists of N-terminal and C-terminal domains, each containing a pair of EF-hand motifs ^9^. The N-terminus of recoverin is acylated by a myristoyl group, which is sequestered in a hydrophobic pocket in calcium-free protein and becomes exposed in response to cooperative binding of Ca^2+^ to EF-hand 3 (high-affinity) and EF-hand 2 (low-affinity) sites ^10,11^. This so-called “Ca²^+^-myristoyl switch” enables the regulatory activity of the protein, as the myristoyl group anchors it to rod outer segment (ROS) membranes, whereas the open hydrophobic pocket forms a binding site for the amphipathic alpha-helix of GRK1 ^5,12,13^. Another important regulatory region of recoverin is the “C-terminal segment”, a variable structural element among NCS proteins that ensures the correct Ca²^+^-sensitivity and architecture of the target-binding site of the protein ^14–16^. Recoverin does not directly affect GRK1 activity, but prevents the C-terminus of rhodopsin (substrate) from reaching the catalytic center of the enzyme ^17^.

Despite the general agreement between the results of structural, functional, and physiological studies of recoverin, no solid explanation has yet been found for the discrepancy between its affinity for Ca²⁺ in vitro (14-19 μM in solution ^14,15,18^) and the physiological range of calcium concentrations (from approximately 500 nM in dark-adapted photoreceptors to 10-50 nM under illumination ^19^), which calls into question the actual ability of recoverin to operate in photoreceptors. Several factors have been proposed to enhance recoverin activity under physiological conditions. First, it is the binding to phospholipid membranes and GRK1, which drives the Ca^2+^-induced structural transitions of Rec ^14,15,18^. Second, it is the presence of high levels of cholesterol and caveolin-1 in membrane microdomains, the binding of which increases the affinity of recoverin for calcium ^20,21^. Third, it is the synergistic action of recoverin with calmodulin, facilitating GRK1 inhibition at physiological calcium levels ^22^. Previous studies indicate that such factors may also include zinc, which binds to both apo and 2Ca^2+^-bound conformers of the protein and alters its structural stability, hydrophobicity, and membrane affinity ^23,24^. These observations are particularly important as zinc is a component of most phototransduction cascade proteins, including rhodopsin (zinc deficiency causes visual impairment) and has recently been recognized as a signaling factor ^25,26^.

Recent investigations have illuminated a more complex regulatory landscape for recoverin and other NCS proteins, indicating that zinc may affect sensitivity of these proteins to redox environments via conservative redox-sensitive cysteine (C39 in recoverin), which may modulate their function ^27–32^. Notably, a disruption of ocular homeostasis of calcium and zinc is associated with retinal neurodegenerative disorders, such as age-related macular degeneration (AMD) and glaucoma ^33–35^. As a protein sensitive to both Ca²⁺ and Zn²⁺, recoverin may respond to abnormal interplay between these signaling metals in photoreceptors, thereby mediating their pathogenic effects on retinal neuron function and survival in human diseases.

Most biochemical and biophysical studies of recoverin have traditionally been conducted using bovine (bRec) and mouse (physiological studies) proteins, while only one work have examined functionally human recoverin (hRec)^36^, despite its obvious significance from a fundamental and biomedical point of view. Although sequence identity between bRec and hRec is high (88%), the differing 24 residues are localized in the regions that may play a crucial role in calcium coordination (EF-hand loops), the Ca^2+^-myristoyl switch mechanism and signaling activity (C-terminal segment), and other properties of the protein (Fig. 1A). Whether hRec functions in a manner that is common to other orthologs or in a specific manner remains unknown. Moreover, the biochemistry of bovine (crepuscular activity type) and mouse (nocturnal activity type) visual systems may differ from that of humans (diurnal lifestyle), which may support unique modes of hRec operation, both in health and disease.

**Figure 1.**
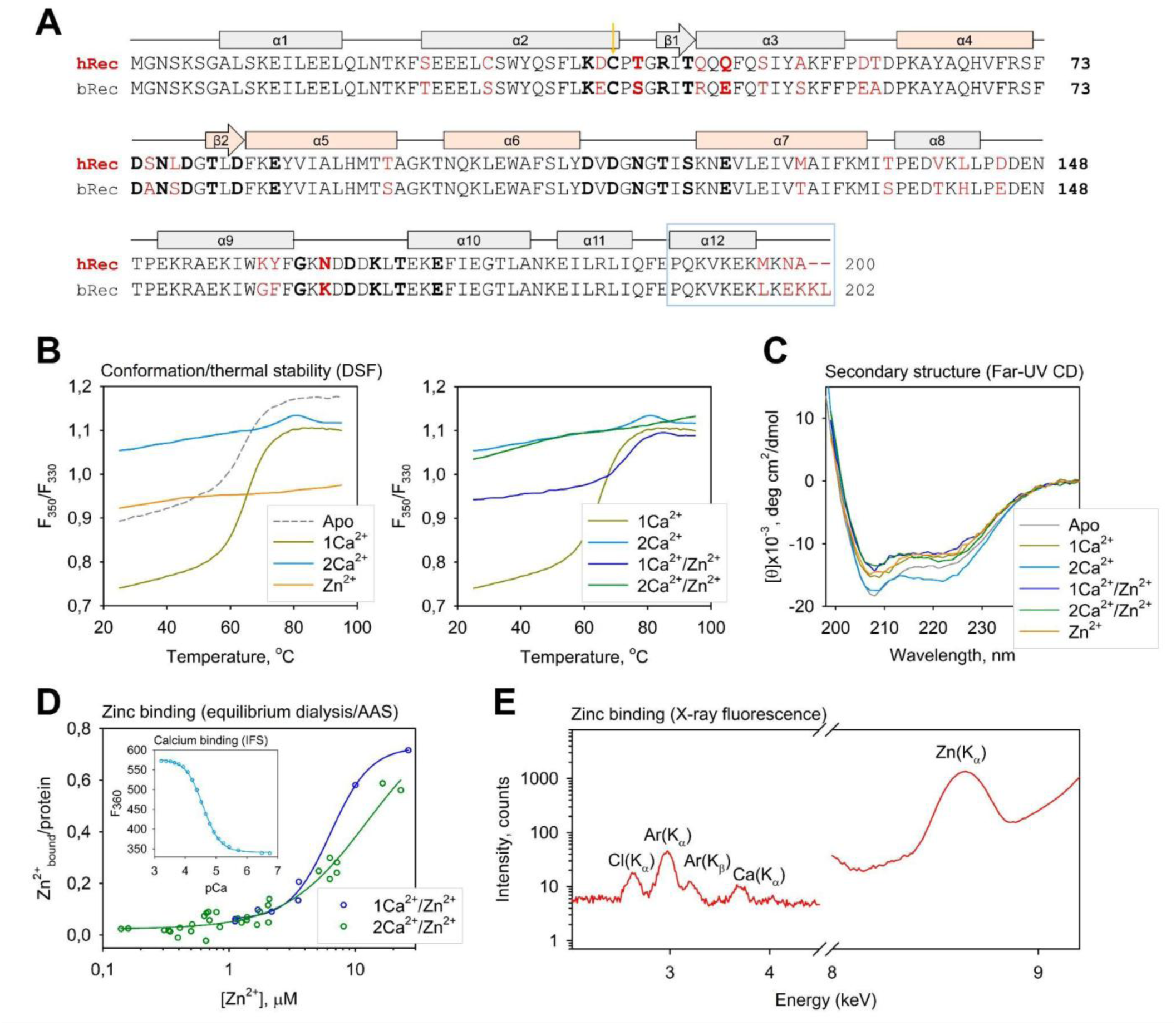
Structural and metal-binding properties of hRec. (**A**) Sequence alignment of hRec and bRec. Non-conservative residues are shown in red. Secondary structure elements are indicated on the top; The amino acids constituting the EF-hand loops are marked in bold, while functional EF-hands are highlighted in rose. Redox-sensitive cysteine (C39) and C-terminal segment are denoted by an arrow and by a blue box, respectively. (**B-C**) Identification of the metal-bound conformers of hRec. NanoDSF thermal unfolding profiles (**B**) and far-UV CD spectra at 25°C (**C**) of hRec forms. (**D-E**) Metal-binding properties of hRec. (**D**) Binding of Zn^2+^ to 1Ca^2+^-bound and 2Ca^2+^-bound hRec according to equilibrium dialysis coupled with AAS. The inset illustrates spectrofluorimetric calcium titration of apo hRec using Ca^2+^-buffer solutions. Solid curves represent the best fits of the data to the 4-parameter Hill equation. (**E**) X-ray fluorescence spectra of the monolayer formed by hRec preincubated with Ca^2+^ and Zn^2+^.

The current study presents for the first time the crystal structure of the 2Ca²⁺-bound form of hRec and provides a comprehensive characterization of its functional properties, revealing the classical structural organization of the protein but highlighting its specific features, such as submicromolar sensitivity to calcium corresponding to the physiological range. Our results also show how the architecture of Ca^2+^-loaded hRec coordinates zinc ions, providing a basis for the regulatory role of Zn^2+^ in photoreceptors. By comparing with the corresponding data obtained for bRec, we identify species-specific differences in the conformational dynamics and functionality of recoverin, providing insight into its evolutionary specialization.

## 2. Results

### 2.1. Metal-bound conformers of human recoverin

To examine the metal-dependent properties of hRec, we prepared apo-form of the recombinant myristoylated protein using bacterial expression, chromatographic purification, and decalcification procedures ^36^. A similarly produced and processed preparation of bRec was obtained for comparative analysis. Both hRec (Fig. S1) and bRec proteins have a purity of >90% and a degree of myristoylation of >95%. The primary screening of metal-bound forms of hRec was performed by monitoring conformation and thermal stability of the protein using nanoDSF and by analyzing its secondary structure using far-UV CD spectroscopy. The nanoDSF study reveals six distinct conformers of the protein, exhibiting unique thermal denaturation profiles (Fig. 1B). In addition to the apo-form, those include the 1Ca^2+^-bound and 2Ca^2+^-bound states, which are achieved in the presence of a single molar calcium excess and saturating concentration of calcium, respectively. All the above-mentioned conformers are able to coordinate zinc, yielding Zn^2+^-bound, 1Ca^2+^/Zn^2+^-bound, and 2Ca^2+^/Zn^2+^-bound forms. The coordination of the first Ca^2+^ (1Ca^2+^-bound form) has a minor effect on the thermal stability of hRec, while coordination of the second Ca^2+^ (2Ca^2+^-bound form) significantly stabilizes the protein, and a similar effect is achieved by Zn^2+^ binding (1Ca^2+^/Zn^2+^-bound form, Table 1). The formation of the described conformers of hRec is generally confirmed by the far-UV CD spectroscopy data (Fig. 1C; Table 1). Each conformer is characterized by a unique ratio of secondary structure elements, although the overall changes caused by metal binding are moderate (Table 1). For comparison, bRec adopts the same conformations and exhibits similar effects related to metal coordination, except that the conformation associated with 1Ca^2+^ exhibits increased thermal stability compared to the apoform, and zinc binding to it does not produce additional effects in this regard (Table S1; Fig. S2A-B).

**Table 1.** Characteristics of various conformers of hRec.

| Structural properties |  |  |  |  |  |  |  |
| --- | --- | --- | --- | --- | --- | --- | --- |
| Thermal stability | Conformer | Apo | 1Ca<sup>2+</sup> | 2Ca<sup>2+</sup> | Zn<sup>2+</sup> | 1Ca<sup>2+</sup>/Zn<sup>2+</sup> | 2Ca<sup>2+</sup>/Zn<sup>2+</sup> |
|  | T<sub>1/2</sub>, °C | 62.3±0.3 | 64.0±0.1 | 75.28±0.14 | n.d. | 70.8±0.4 | n.d. |
| Secondary structure content, % | A | 53.0±1.2 | 47.3±1.2 | 55.4±1.2 | 45.8±1.2 | 45.75±1.2 | 48.6±1.2 |
|  | B | 6.4±0.2 | 8.3±0.2 | 6.1±0.2 | 8.8±0.2 | 9.9±0.2 | 8.3±0.2 |
|  | Turns | 15.2±0.3 | 17.4±0.3 | 14.2±0.3 | 18.5±0.3 | 18.2±0.3 | 16.7±0.3 |
|  | Disordered | 25.7±0.8 | 26.8±0.8 | 23.9±0.8 | 27.0±0.8 | 25.6±0.8 | 25.5±0.8 |
| Metal-binding properties |  |  |  |  |  |  |  |
| Calcium binding | K<sub>D</sub>, μM | 26.8±0.4 |  |  |  |  |  |
|  | N | 1.56±0.03 |  |  |  |  |  |
| Zinc binding | Conformer | 1Ca<sup>2+</sup> |  |  | 2Ca<sup>2+</sup> |  |  |
|  | K<sub>D</sub>, μM | 6.4±0.6 |  |  | 12±5 |  |  |
|  | N | 2.7±0.4 |  |  | 1.4±0.3 |  |  |
| Functional properties |  |  |  |  |  |  |  |
| ROS membrane binding | EC<sub>50</sub>, [Ca<sup>2+</sup>] | 0.22±0.11 |  |  |  |  |  |
|  | N | 1.1±0.5 |  |  |  |  |  |
| GRK1 binding | Conformer | Ca<sup>2+</sup> |  |  | Ca<sup>2+</sup>/Zn<sup>2+</sup> |  |  |
|  | K<sub>D</sub>, μM | 17±3 |  |  | 16±3 |  |  |

n.d., not determined

### 2.2. Metal-binding properties of human recoverin

In the next step, we determined the affinities of hRec conformers for calcium and zinc. Calcium binding was monitored by changes in its intrinsic (tryptophan) fluorescence (IFS) using Ca^2+^ buffers (Fig. 1D, inset) ^36^. Fitting the data to the Hill model reveals cooperative binding with a KD of 26.8 μM (Table 1), which is similar to the corresponding parameters of the bovine orthologue (16.7 μM; Table S1). The parameters of zinc binding were measured by atomic absorption spectroscopy (AAS) coupled with equilibrium dialysis (Fig. 1D), focusing on the 1Ca^2+^/Zn^2+^-bound and 2Ca^2+^/Zn^2+^-bound conformers, which represent the novel and physiologically most intriguing forms of the protein. The affinity for zinc of 1Ca^2+^-bound hRec (K_D_ = 6.4 μM) is higher than that of 2Ca^2+^-bound hRec (K_D_ = 12 μM) (Table 1) and generally corresponds to the previously determined value for 2Ca^2+^-bound bRec (K_D_ = 2.7-7.1 μM ^23,24^.

To verify the formation of these conformers, particularly the incorporation of Zn^2+^ into calcium-saturated hRec, we used synchrotron-induced X-ray fluorescence analysis of protein films on the surface of a liquid, which allows reliable detection of trace elements in a protein sample ^37^. The study reveals a pronounced zinc peak in the monolayer formed by hRec preincubated with Ca^2+^ and Zn^2+^ (Fig. 1E), directly demonstrating the ability of the Ca^2+^-loaded hRec to bind zinc. Similar observations were made for bRec (Fig. S2D).

### 2.3. Metal-dependent functional properties of human recoverin

To assess the effects of the Ca^2+^/Zn^2+^ binding on physiological activity of hRec, we next analyzed Ca^2+^-myristoyl switch-related functional properties of the protein, specifically its interaction with ROS membranes and GRK1. Since the components of functional tests may include zinc chelators (e.g., membrane phospholipids), all experiments involving zinc were performed at its saturating concentrations. Therefore, the analysis was limited to Zn^2+^-bound and 2Ca^2+^/Zn^2+^-bound forms of hRec. As shown by the equilibrium centrifugation assay, hRec binds to native ROS membranes in a Ca^2+^-dependent manner (Fig. 2A). Notably, calcium titration revealed half-maximal binding at 0.22 μM [Ca^2+^]_free_, which is 6-fold lower than the corresponding value for bRec (1.3 μM, Table S1) and 122-fold lower than the dissociation constant of the 2Ca^2+^-hRec in aqueous solution (26.8 μM, Table 1). The incorporation of zinc into the Ca^2+^-loaded hRec (2Ca^2+^/Zn^2+^-bound form) augmented its association with ROS membranes, but the effect is insignificant (Fig. 2A, inset).

**Figure 2.**
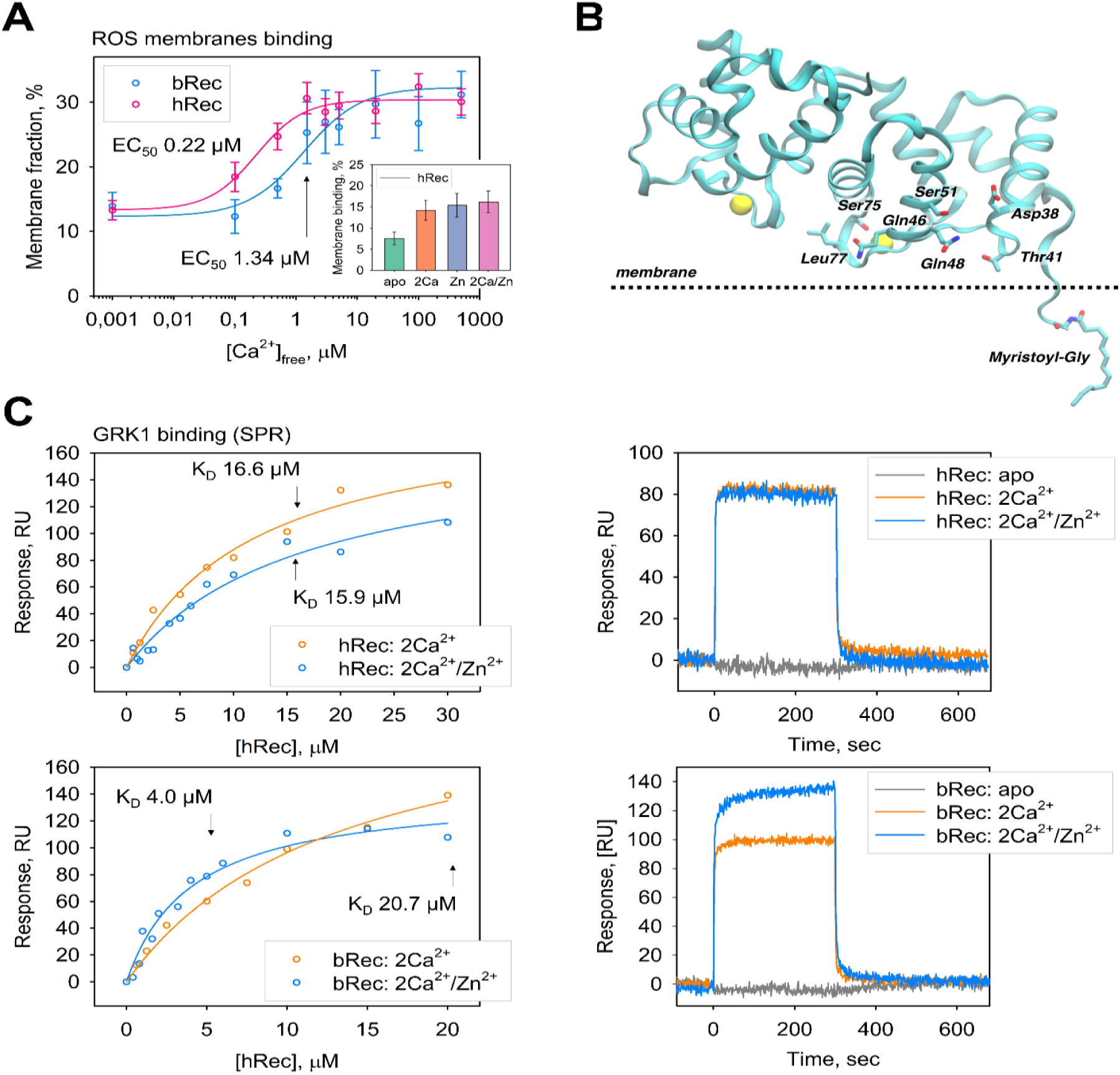
Functional properties of hRec. (**A-B**) Membrane association of hRec and bRec. (**A**) Weight fractions of ROS membrane-bound recoverins at various concentrations of free calcium, controlled using Ca^2+^-buffer solutions, according to the equilibrium centrifugation assay. The inset shows the maximum fractions of the membrane-associated forms of apo-, 2Ca^2+^-bound, 2Ca^2+^/Zn^2+^-bound, and Zn^2+^-bound conformers of hRec. (**B**) MD simulation of the interaction of hRec with ROS membranes. The residues forming the protein-membrane interface and the myristoyl group embedded in the membrane are labeled. Yellow spheres represent Ca^2+^ in EF hands 2 and 3. The analysis is performed using the crystal structure of 2Ca^2+^-bound hRec resolved in this study (see below). (**C**) Binding of hRec and bRec to GRK1. Steady-state analysis of the binding of recoverins to GST-fused GRK1 fragment 1-25, performed using SPR technique. The amplitudes of the equilibrium binding signals are shown as a function of hRec concentration. Solid curves represent best fits of the data to the Langmuir equation. Representative SPR sensorgrams showing GRK1 binding by apo-, 2Ca^2+^-bound and 2Ca^2+^/Zn^2+^-bound conformers of hRec/bRec at 10 μM concentration are shown on the right.

The interaction of hRec conformers with GRK1 was monitored by the SPR biosensor technique. Glutathione S-transferase (GST)-fused GRK1 fragment 1-25, which was previously found to be involved in complexation with recoverin ^5^, was immobilized on the sensor chip, while hRec forms were supplied in the analyte phase (Fig. 2C). The dissociation constant of the complex of GRK1 with Ca^2+^-loaded hRec was determined as 17 μM (Table 1), which is comparable to the corresponding value for the bovine protein (20 μM - Table S1). Interestingly, Ca^2+^/Zn^2+^-bound hRec showed only a negligible change in affinity to GRK1 compared to the Ca^2+^-loaded conformer, whereas in the case of bRec the corresponding increase was 5-fold (Fig. 2C).

### 2.4. Crystal structure of 2Ca^2+^-bound and 1Ca^2+^/2Zn^2+^-bound forms of human recoverin

To gain an insight into the structural basis of Ca²⁺/Zn²⁺-dependent hRec activity, we examined the tertiary structure of the corresponding forms of the non-myristoylated protein using X-ray crystallography (the myristoylated form is highly resistant to crystallization, and no crystal structure of this form has been reported to date). The conditions for hRec crystallization were chosen by varying both pH and Ca²⁺ concentration in the presence of ZnCl₂. As a result, crystals ranging in size from 50 μm to 300 μm belonging to the I4 space group were obtained (Fig. S4). X-ray diffraction analysis of the crystals revealed two unique conformers of hRec coordinating either two calcium ions (2Ca²⁺-bound, G7–Q191) or one calcium ion and two zinc ions (1Ca²⁺/2Zn²⁺-bound, A8–P190) resolved at 1.60 Å and 1.85 Å, respectively (Table 2). Both structures had a characteristic recoverin fold consisting of two domains (residues 7-91, N-terminal, and 100-191, C-terminal) connected by a flexible linker (residues 92-99), with each domain containing a pair of EF-hand motifs (Fig. 3A). The N-terminal end residues were not modeled due to their intrinsic flexibility, consistent with previous bRec structures (PDB 1JSA, 4M2Q, 4M2P, 4MLW, 4YI8, 4YI9, 4M2O, 2HET). The C-terminal portion of the protein, previously determined as the “C-terminal segment” ^14,15,38,39^, was also unresolved in our models. In general, the structure of hRec was similar to reported bRec crystal structures, with a maximum RMSD Cα-Cα of approximately 1.2 Å with the E153A mutant of bRec (PDB 4YI8). The key novel findings were the presence of calcium in both EF-hands of the wild-type protein and the discovery of zinc sites, which we observed for the first time. The RMSD Cα-Cα between 2Ca²⁺-bound and 1Ca²⁺/2Zn²^+^-bound forms was determined as 0.22 Å.

**Figure 3.**
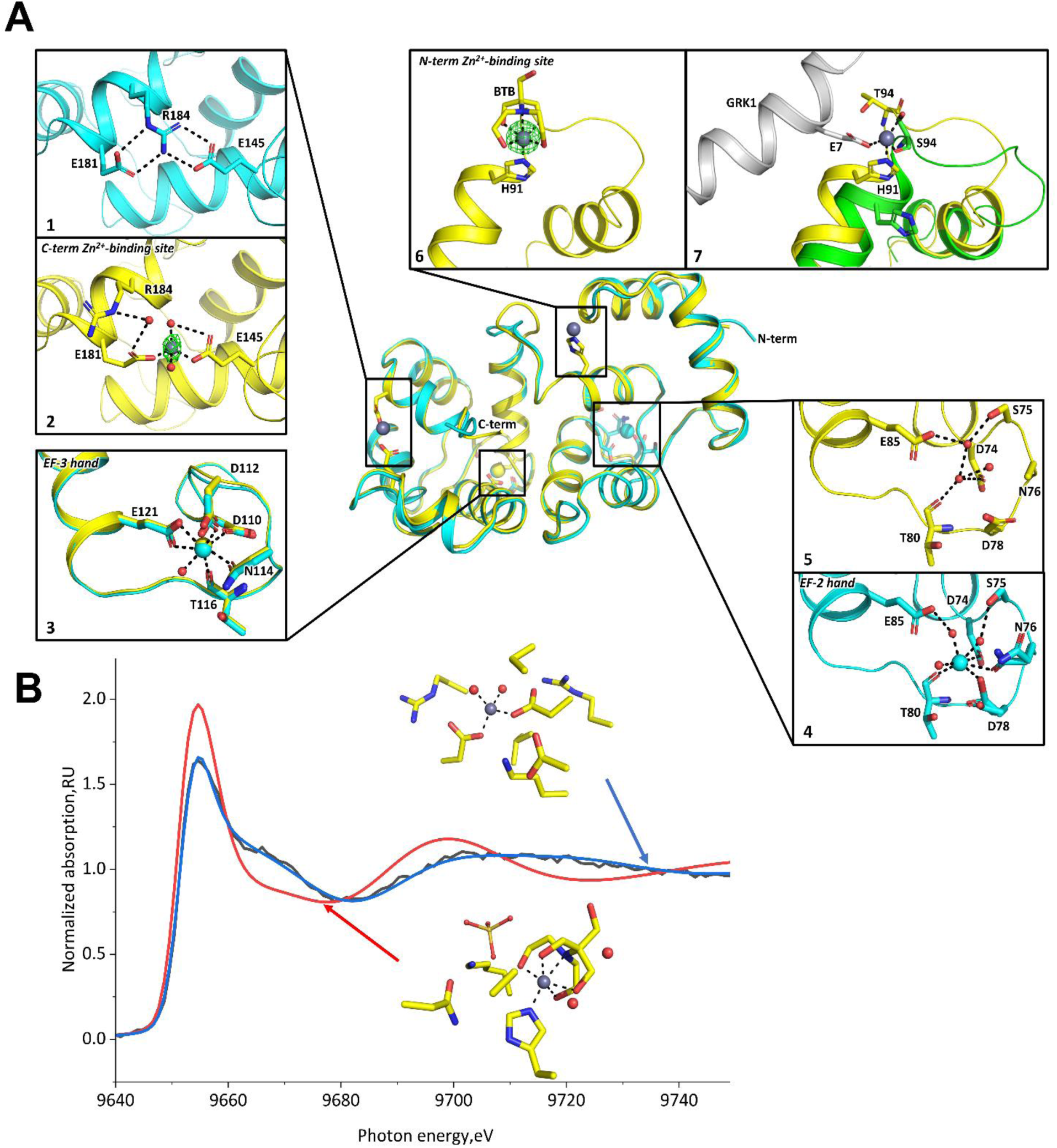
Molecular organization of hRec. (**A**) Crystal structure of 2Ca^2+^-bound (cyan) and 1Ca^2+^/2Zn^2+^-bound (yellow) forms. The spheres represent Ca^2+^ (yellow/cyan), Zn^2+^ (gray), and water (red). The residues involved in the metal coordination are labeled in N-terminal (box 6) and C-terminal (boxes 1-2) Zn^2+^-binding sites, as well as Ca^2+^-binding sites in EF-hand 2 (boxes 4-5) and EF-hand 3 (box 3). The anomalous difference map (green mesh) surrounding the zn^2+^ ions is contoured at 8σ. (Box 7) Expected position of the N-terminal Zn^2+^, mediating the hRec (yellow)/bRec (green) and GRK1 (gray; PDB 2I94) complex. (**B**) Experimental (black) and theoretical four-coordinate (blue) and six-coordinate (red) XANES spectra at the zinc K-edge.

**Table 2.**
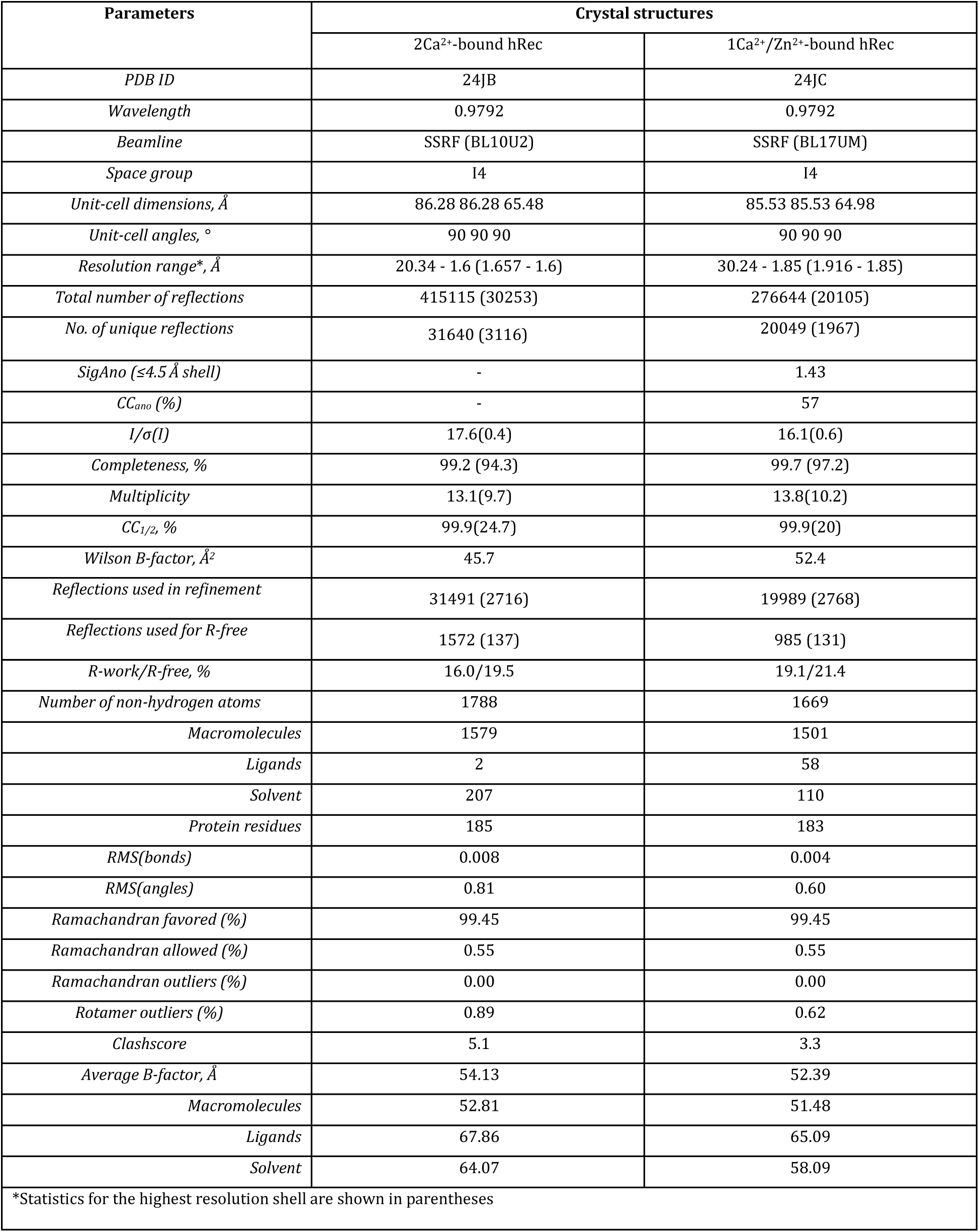
X-ray diffraction data collection and refinement statistics.

In 2Ca²⁺-bound hRec, both sites exhibit classical EF-hand geometry, with Ca²⁺ coordinated by seven oxygen atoms. In EF-hand 3, it is bound in a pentagonal bipyramidal configuration by side chain oxygen atoms of D110, D112, N114, and E121 (bidentate ligand), as well as the main chain carbonyl oxygen of T116 and an axial water molecule (Fig. 3A, *box 3*). In EF-hand 2, Ca²⁺ is coordinated by side chain oxygens of D74, N76, D78, main chain carbonyl of T80, and three water molecules, adopting a distorted pentagonal bipyramidal (Fig. 3A, *box 4*). Partial occupancy (85%) of the Ca²⁺ ion and alternative conformations of N76 and D78 were observed, suggesting reduced stability compared to EF-hand 3.

In the 1Ca^2+^/2Zn^2+^-bound hRec, EF-hand 3 exhibits a similar Ca^2+^-loaded conformation, while EF-hand 2 remains occupied by water molecules, leading to disorder in the N76 sidechain (unresolved) (Fig. 3A, *box 5*). Data collected at a wavelength of 0.979 Å showed strong anomalous difference peaks (28σ and 11σ) corresponding to zinc ions in the N-terminal (Fig. 3A, *boxes 6 and 7*) and C-terminal domains outside the functional EF-hands (Fig. 3A, *box 2*). Using the single-wavelength anomalous dispersion data, we refined two zinc-binding sites within the asymmetric unit with occupancies of 100% (N-terminal domain) and 55% (C-terminal domain). The anisotropic displacement parameters (ADPs) are closely aligned with the neighboring atoms. The first (N-terminal) site is located at H91, where N𝜀 atom of the imidazole ring forms a hydrogen bond with a coordinating water molecule, while N𝛿 directly coordinates the zinc (Fig. 3A, *box 6*). The other coordination positions are provided by buffer component Bis-Tris, BTB (2-[bis-(2-hydroxy-ethyl)-amino]-2-hydroxymethy-propane-1,3-diol), completing the octahedral geometry. At the second (C-terminal) site, zinc exhibits a classic tetrahedral coordination geometry with carboxylate oxygen atoms from E145 and E181, and two water molecules (Fig. 3A, *box 2*).

### 2.5. X-ray absorption spectroscopy study of zinc environment in human recoverin

To validate the zinc binding site 2 identified in the crystal structure of hRec, we employed X-ray absorption spectroscopy (XAS) targeting the zinc K-edge. This technique allowed us to investigate the local coordination environment by analyzing X-ray absorption near-edge structure (XANES), which provides detailed information about the short-range electronic and geometric structure surrounding the bound zinc atom. The K-edge XANES spectrum is derived from dipole-allowed transitions of zinc 1s core electrons to unoccupied 4p states, which are strongly influenced by the chemical environment via hybridization with surrounding ligand atoms. To confirm the Zn^2+^ coordination geometry, we computed theoretical XANES spectra for the coordination numbers of four and six, using the QM/MM-optimized geometries of the zinc-binding site (Fig. 3B). The distinct tetrahedral and octahedral coordination geometry arrangements around the zinc atom modify configurations of the unoccupied 4p states, leading to the characteristic spectral features. Specifically, the tetrahedral environment results in reduced local symmetry compared to an octahedral arrangement. This reduction in the symmetry manifests in the XANES spectrum as a splitting of the primary absorption peak just beyond the edge. Comparison of the theoretical XANES spectra with experimental data for hRec revealed that the key spectral characteristics, including peak positions and absorption intensities, are well reproduced by the tetrahedral coordination model, whereas the octahedral coordination does not match the observed data. These results demonstrate that zinc is bound to hRec under solution conditions in tetrahedral coordination, in accordance with the C-terminal Zn^2+^-binding site observed in the crystal structure (Fig. 3A, box 2).

### 2.6. Molecular dynamics simulations of 2Ca^2+^-loaded human recoverin bound to lipid membranes

The key distinguishing feature of hRec compared to its bovine ortholog is its enhanced association with ROS membranes, reflected in a 6-fold lower calcium concentration required for the binding (Table 1 and Table S1). To elucidate the structural determinants underlying this difference, we used the crystal structure of 2Ca^2+^-bound hRec resolved in this study to perform MD simulations in a membrane environment consisting of POPE: POPG bilayer. The initial position of the recoverin molecule was chosen based on previous data ^12,13^, showing that its myristoylated N-terminus engages with the membrane surface, while the C-terminal domain remains oriented away from the lipid bilayer (Fig. 2B). The system was simulated for 0.5 𝜇s without restraints to refine the membrane-bound protein conformation and quantify protein–membrane interactions. Comparison of per-residue solvent-accessible surface area (SASA) values in the presence and absence of the membrane enabled identification of the residues that directly contact the bilayer (Table S2). Among these membrane-contacting residues, we next focused on those that differ between bovine and human recoverin sequences. This analysis yielded a short list of the substitutions that may contribute to the increased membrane affinity of the human variant: T41S, L77S, S75A, Q46R, Q48E, D38E, and S51T. To evaluate the energetic impact of each substitution, we performed thermodynamic integration (TI) free energy calculations: for each mutation, alchemical transformations were conducted in both membrane-bound and membrane-free states, enabling determination of ΔΔG values through the thermodynamic cycle shown in Fig. S6. The resulting ΔΔG values provide a quantitative estimate of how each substitution influences the membrane affinity of recoverin (Table 3). The most energetically favorable substitution identified was replacement of the negatively charged E48 in bRec with Q, which enhances membrane binding of hRec by approximately -22 kJ mol⁻¹. This effect is followed by the substitution of the positively charged R46 by Q (-11 kJ mol⁻¹) and the more modest impact of the E38D mutation (-7 kJ mol⁻¹). In contrast, the A75S replacement impedes membrane binding by +11 kJ mol⁻¹. When combined, the cumulative effect of the seven substitutions on the free energy of membrane binding amounts to -30 kJ mol⁻¹, consistent with the experimentally observed increased binding level in the case of hRec. The high energetic contribution associated with the E48 residue in bRec may reflect not only the release of electrostatic repulsion from anionic POPG lipids in the model ROS membrane but, more generally, the high energetic cost of the removal of the buried charge at the interface between two low-dielectric media, i.e., protein and membrane ^40,41^, in line with a similar effect of the R46 residue on the membrane binding. Conversely, the A75S substitution introduces a polar side chain into the interfacial region, thereby increasing the desolvation penalty and preventing membrane association.

**Table 3.** Change in the free energy of membrane binding of recoverin induced by point mutations distinguishing hRec and bRec.

| Human | Bovine | $\Delta SASA, \text{\AA}^2$ | $\Delta\Delta G^{\text{human-bovine}}, \text{kJ/mol}$ |
| --- | --- | --- | --- |
| THR41 | SER | 70.9 | -2.1±1.3 |
| LEU77 | SER | 34.0 | 0.6±1.5 |
| SER75 | ALA | 17.1 | 10.7±0.7 |
| GLN46 | ARG | 15.1 | -10.6±1.8 |
| GLN48 | GLU | 14.5 | -21.7±1.3 |
| ASP38 | GLU | 3.8 | -7.4±1.8 |
| SER51 | THR | 3.2 | 0.1±0.21 |

## 3. Discussion

This study presents the first comprehensive structural and functional characterization of hRec, refining the activity of this key photoreceptor protein in humans. In general, hRec behaves similarly to the well-characterized bovine ortholog, forming Ca^2+^-bound, Zn^2+^-bound, and Ca^2+^/Zn^2+^-bound conformers and undergoing a Ca^2+^-myristoyl switch governing its interaction with photoreceptor membranes and GRK1. At the same time, it demonstrates a number of specific features. First, the effect of the increased sensitivity to calcium in the presence of phospholipid membranes in the case of hRec is much more pronounced (122-fold) compared to bRec (13-fold). Second, in hRec, calcium binding leads to the formation of a specific zinc site(s) as a result of Ca^2+^-induced conformational change, the filling of which stabilizes the protein, at least in the case of 1Ca^2+^-bound form. Third, hRec and bRec show different thermal stability of Ca^2+^/Zn^2+^-bound conformers and their behavior in binding with GRK1. Thus, zinc binding stabilizes the active (open) conformation of 1Ca^2+^-bound hRec, but only in bRec does zinc enhance its affinity for GRK1, since the latter likely directly participates in the coordination of the metal (see below).

The observed phenomenon of membrane-induced enhancement of hRec sensitivity to Ca^2+^ is consistent with the cellular conditions of its functioning and resolves the problem of its low calcium affinity in aqueous solutions. Thus, although the dissociation constant of the hRec complex with calcium is relatively high (26.8 µM), the determined half-maximal Ca^2+^ concentration for hRec binding to photoreceptor membranes (220 nM) is much lower and falls into the region of light-induced changes in free calcium concentration in photoreceptors (from ∼500 nM in the dark to 10-50 nM upon illumination) ^19^. This differs strikingly from the estimates for bRec (1.3 μM; this study), and for the first time demonstrates the possibility of operation of the Ca²⁺-myristoyl switch of recoverin under physiologically relevant free calcium conditions.

Zinc seems to be another factor supporting the physiological activity of recoverin, as previously reported for rhodopsin, PDE6, and other components of the visual cascade ^42,43^. Earlier, we suggested that bRec may act as a physiological Ca^2+^/Zn^2+^ sensor protein ^25,26,44^, but whether zinc binding is characteristic of normal retinal physiology or occurs only under pathological conditions remains unclear. In general, being a component of visual cascade proteins, zinc plays a key role in phototransduction, and its deficiency causes night blindness^45^. The total zinc concentration in the retina is high (hundreds of micromoles), and a significant portion may exist as a chelatable (“labile” or “loosely bound”) Zn^2+^, apparently possessing signaling function ^25,26,44^. In bovine retina, the highest zinc concentration is found in the outer segments and synaptic terminals of photoreceptors ^46^. The release of necessary chelatable zinc can follow intracellular acidification at high calcium in the dark ^47,48^. Under these conditions, zinc will thereby colocalize with 2Ca^2+^-bound Rec associated with ROS membranes and may be coordinated by it, as shown for rhodopsin ^42,49,50^. The formation of 2Ca^2+^/Zn^2+^-bound bRec, exhibiting enhanced affinity towards GRK1 (see Fig. 2C), might prolong the lifetime of photoexcited rhodopsin and favor high sensitivity of bovine retinal rods to light. In the case of hRec, no such effect is observed, which can underlie poorer night vision of humans compared to cattle (diurnal versus crepuscular behavior) ^51,52^. An analogous study comparing bovine and human rhodopsins revealed similarities in the architecture of the photoexcited G-protein-activating conformer, but differences in the inactive receptor ^53^. Thus, alterations in rhodopsin activation and desensitization (modulated by recoverin) in host species can be related to their light activity patterns, e.g., diurnality or crepuscularity. In the case of hRec, the conditions for *in vivo* generation of its 1Ca^2+^/Zn^2+^-bound form may arise under weak/unsaturated illumination, which leads to a moderate decrease in calcium concentration ^54^, sufficient for clearance of the low-affinity EF-hand 2. In general, 1Ca^2+^/Zn^2+^-bound hRec appears to be the most favorable form as it is characterized by elevated thermostability (Table 1) and is formed during crystallization at the saturating calcium and zinc levels. By stabilizing 1Ca^2+^-hRec in the open state (see below), zinc may prolong its membrane and GRK1 binding, thereby supporting the activity of the protein under these conditions.

Despite the different but generally beneficial effects of zinc on Ca^2+^-saturated hRec and bRec forms, it tends to destabilize the Ca^2+^-free forms of both orthologs, which are formed at saturating light intensities. Under these conditions, Zn^2+^-bound hRec/bRec can accumulate not only in the outer but also in the inner segments of photoreceptors, where both substances are being translocated ^55–57^. Moreover, severe oxidative stress, accompanying light-induced retinal degeneration, results in the accumulation of chelatable zinc in these compartments ^58^, thereby promoting the generation of such forms. Consistently, elevated zinc levels were found directly in the retina during acute degeneration, apparently due to its redistribution from the RPE ^33^. Zinc is known to stimulate thiol oxidation of bRec ^28^, which suppresses functionality of the protein, including its ability to inhibit GRK1 ^27,29,30,59^. Accordingly, it was found that light-induced retinal degeneration is associated with prolonged phosphorylation of rhodopsin by GRK1 ^60^. Overall, zinc appears to have a dual effect on recoverin activity, namely enhancing it under normal conditions, which contributes to the light sensitivity of bovine retinal rods, but weakens it due to protein thiol oxidation under conditions of oxidative stress associated with degenerative retinal diseases such as AMD.

The structural features underlying the specific behavior of hRec were determined by analyzing its molecular organization using X-ray crystal structure complemented with XANES and MD analysis. hRec has 88% sequence identity with bRec (Fig. 1A), but in certain structural aspects, these proteins are quite distinct. Despite that hRec crystallized in a similar tetragonal space group as bRec, its crystals exhibit slightly larger unit cell dimensions (c ≈ 65 Å versus 58–60 Å) and a higher solvent content and Matthews coefficient (Table S3), indicating a more hydrated lattice and greater flexibility of the human protein (Fig. S5). This crystal cell swelling in hRec likely results from the destabilization of the E59–E181’ and E143–T45’ (the prime denotes a symmetry-related molecule) hydrogen bonds that anchor the bRec crystal packing (Fig. S5C); in hRec, these interactions are disrupted by the mutation of glutamate to shorter aspartate residues. Additionally, the G157K substitution in hRec introduces a bulky sidechain (see Fig. S5D), which disrupted the direct crystal contact (G157-S41’) in bRec (Fig. S5C), further contributing to the lattice expansion.

The first key structural difference between hRec and bRec concerns the C-terminal segment, a variable regulatory element among NCS proteins (boxed in Fig. 1A). In bRec, this region is stabilized by a hydrophobic cluster formed by residues I133, L141, V193, and L197, as well as by electrostatic interactions between the positively charged residues K194, K198, and H140 and the negatively charged residues E136 and D137 ^14^. In hRec, both of these stabilizing clusters are disrupted by the L197M and H140L substitutions, correspondingly. As a result, the C-terminal segment appears destabilized and becomes unresolved in the hRec structure beyond Q191, likely owing to increased flexibility caused by the loss of these anchoring interactions.

The second key difference concerns the filling of the functional EF-hands. In all previous crystallographic studies of wild-type bRec, calcium was detected only in EF-hand 3 (PDB 1OMR, 4M2Q, 4MLW) ^32,61^. The coordination of Ca^2+^ in EF-hand 2 remained elusive due to intermolecular crystal contact between T80 and E153’ artificially preventing Ca²⁺ coordination (Fig. S5B), and this limitation was overcome by introducing the E153A mutation (PDB 4YI8) ^62,63^. In our hRec structure, E153’ lacks contact with T80 due to the crystal cell swelling, which facilitates Ca²⁺ coordination in EF-hand 2. Instead, E153 forms a salt bridge with K119 (Fig. S5B), likely enhancing the protein’s overall stability and function. Thus, we report here the first crystal structure of wild-type recoverin with two Ca²⁺ ions bound at EF-hands 2 and 3. It should be added that the side chain of E85, known to be implicated in calcium coordination in bRec (PDB 4YI8 and 1JSA) ^62,63^, did not show direct interaction with Ca²⁺ in our hRec structure; instead, the interaction is mediated by a water molecule (Fig. 3A, box 4).

Finally, the third key difference between hRec and bRec concerns the membrane-binding interface. As detailed in Section 2.6, specific substitutions at the protein surface account for the notably higher membrane affinity of hRec compared to bRec. The MD simulations identify the E48Q substitution in hRec as the most critical factor: the presence of a negatively charged glutamate at this position in bRec introduces a significant +22 kJ/mol destabilization penalty relative to the glutamine in hRec (Table 3). This effect likely stems from both electrostatic repulsion with anionic lipids and the high energetic cost of burying a charge at the low-dielectric protein-membrane interface.

It should be noted that our work is the first to directly show the structure of zinc-binding sites in the NCS protein family, although the zinc sensitivity in this family has been extensively characterized ^23,24,64^. Our X-ray crystallographic analysis identified two zinc ions under crystallization conditions: a four-coordinate C-terminal site and a six-coordinate N-terminal site (in part, formed due to the buffer molecule). However, XANES spectroscopy showed that only the C-terminal is occupied in solution; this site is formed by the side chains of E145 and E181, along with two coordinating water molecules (Fig. 3A, box 2). However, in the 2Ca^2+^-bound structure, this site is occupied by the canonical internal salt bridge (Fig. 3A, box 1). Thus, in addition to calcium-free and 2Ca^2+^-bound forms, the Ca^2+^/Zn^2+^-bound form may be a physiologically relevant variant of hRec. It represents a stable open conformer (Table 1), in which zinc binding stabilizes a functionally important region reinforced by a hydrogen bond between carboxylate E181 and the guanidine group of R184 via a water molecule. The N-terminal zinc-binding site at H91 is formed with the participation of Bis-Tris molecules, suggesting its partially artificial nature (Fig. 3A, box 6). However, under physiological conditions, the Bis-tris positions may be occupied by other endogenous ligands, such as natural buffers or water molecules, thereby restoring this site. For instance, zinc coordination in the N-terminal site can be complemented in complex with GRK1. Indeed, alignment of the structure of our 1Ca^2+^/2Zn^2+^-bound hRec and the bRec–RK25 complex (PDB 2I94) ^5^ revealed that E7 residue of GRK1 approaches the zinc binding site and could directly participate in zinc coordination, replacing Bis-tris as a ligand (Fig. 3A, box 7). Other coordinating atoms in this case could be the carbonyl and hydroxyl oxygens from S94. Such a configuration would allow zinc to act as a structural bridge between bRec and GRK1, thereby enhancing the stability and affinity of this complex (see Table 1). In human protein, the same effect will be weaker owing to the replacement of S94 with the bulkier T94, although zinc can still interact with His91 of recoverin and E7 of GRK1, as well as with water molecules. These observations explain the lower stability of the Zn^2+^-hRec complex with GRK1 (Table 1), which may represent a key physiological difference between human and bovine orthologs.

## 4. Conclusion

We report the first crystal structures of 2Ca²⁺-bound and 1Ca²⁺/2Zn²^+^-bound hRec, revealing the structural basis for its calcium- and zinc-mediated regulation. Our work is the first to directly show the structure of zinc-binding sites in the NCS family, although zinc sensitivity in the NCS family has been extensively characterized. Compared to bRec models, hRec shows species-specific conformational differences, offering insights into their evolutionary specialization. Functionally, these differences are related to calcium sensitivity of membrane association, responses to zinc and GRK1 binding, and may be associated with the specific conditions in the human retina. We note that bRec cannot be considered as an accurate template for hRec, and their structural and functional differences must be taken into account. This may lead to a revision of some commonly accepted concepts of rhodopsin desensitization and general mechanisms of vision in humans. In addition to hRec specificity, our findings highlight the general importance of recoverin as a key node linking calcium and zinc signaling pathways in mammalian photoreceptors under normal and pathological (oxidative stress) conditions.

## 5. Materials and Methods

### 5.1. Protein Expression and Purification

Recombinant forms of hRec and bRec were obtained according to the previously developed procedure ^36^. Briefly, the proteins were expressed in *E. coli* strains pET15b rec/pBB131/BL21(DE3)RILC+ (myristoylated protein) and pET15b rec/BL21(DE3)RILC+ (non-myristoylated protein). To obtain myristylated protein, myristic acid (0.2 g/L) was added to the growth medium immediately after the induction. The cells were cultured for 22 h at 30°C, harvested by centrifugation at 5,000×g for 15 min at 4°C, resuspended in lysis buffer (50 mM Tris-HCl, 100 mM NaCl, 1 mM DTT, 1 mM EDTA, pH 8.0), and lysed using RETSCH (Haan, Germany) mixer mills. The lysate was centrifuged at 25,000×g for 40 min at 4°C. The supernatant was adjusted to 10 mM CaCl₂, filtered through cotton wool, and applied to a phenyl-Sepharose column equilibrated with buffer A (20 mM Tris-HCl, 1 mM DTT, 10 mM CaCl₂, pH 8.0). The column was washed with buffer A, followed by buffer A without CaCl₂, and recoverin was eluted using buffer A containing 10 mM EDTA. Removal of calcium and metal chelate complexes from the obtained protein preparations was carried out by gel filtration as described in ^65^. Purified recoverin samples were concentrated using Amicon Ultra-15 centrifuge filters (Merck Millipore Ltd., Darmstadt, Germany) at 7,000×g for 20 min at 4°C, and stored at -70°C. The molecular weight and degree of myristoylation of the obtained recombinant proteins were verified by liquid chromatography–mass spectrometry. The samples were applied to a Jupiter C18 column (Phenomenex, Torrance, CA, USA) pre-equilibrated with 0.05% trifluoroacetic acid and 10% acetonitrile, eluted with an acetonitrile gradient and analyzed using LCMS-2010EV mass spectrometer (Shimadzu, Kyoto, Japan). Theoretical molecular mass calculations and electrospray ionization mass spectra deconvolution were performed using GPMAW ^66^ and MagTran software ^67^, respectively. Recoverin concentrations were determined spectrophotometrically using a molar extinction coefficient at 280 nm of 25,440 M⁻¹ cm⁻¹ and 24,075 M⁻¹ cm⁻¹ for human and bovine proteins, respectively.

Genetic constructs encoding fragment 1-25 of rhodopsin kinase (GRK1), fused with glutathione-S-transferase (GST), and GST, inserted into the pGEX-5x-1 vector, were expressed in the *E. coli* BL21-CodonPlus(DE3) strain and the respective proteins were purified by affinity chromatography on Glutathione Sepharose 4B (Cytiva, Marlborough, MA, USA) as described previously ^17,22.^

### 5.2. Differential Scanning Fluorimetry (nanoDSF)

Primary screening for metal-bound conformers of hRec and bRec was performed using nanoDSF with Prometheus Panta (NanoTemper Technologies, Munich, Germany) and PSA-16 (Biowe, China). Prometheus Standard capillaries (NanoTemper Technologies) were loaded with 10 μM hRec in 20 mM Tris-HCl buffer (pH 8.0), 150 mM NaCl, containing one of the following components: 1 mM EDTA (apo form), 10 µM CaCl_2_ (1Ca^2+^-bound form), 1 mM CaCl_2_ (2Ca^2+^-bound form), 10 µM CaCl_2_/50 µM ZnCl_2_ (1Ca^2+^/Zn^2+^-bound form), or 1 mM CaCl_2_/50 µM ZnCl_2_ (2Ca^2+^/Zn^2+^-bound form). The samples were heated from 25 °C to 95 °C at a rate of 0.5 °C/min. The melting temperature at the midpoint of the thermal denaturation transition (T_m_) for each sample was calculated from the temperature dependence of the first derivative of the ratio of tryptophan fluorescence intensities at 350 nm and 330 nm (F₃₅₀/F₃₃₀). The excitation wavelength was 280 nm. The data obtained were analyzed using PR.Panta Analysis software (NanoTemper Technologies).

### 5.3. Circular Dichroism (CD) spectroscopy

Circular dichroism (CD) studies of myristoylated forms of hRec and bRec conformers were carried out at 25°C with a J-810 spectropolarimeter (JASCO Inc., Tokyo, Japan), equipped with a Peltier-controlled cell holder. The instrument was calibrated with an aqueous solution of d-10-camphorsulfonic acid according to the manufacturer’s instructions. The cell compartment was purged with nitrogen (dew-point of –40°C). The quartz cell with a pathlength of 1.00 mm was used for far-UV measurements. Protein concentration was 1.7 – 2.8 µM. Buffer conditions: 20 мМ Tricine-KOH pH 7.3, 50 мМ KCl, 20 μМ DTT, 100 μM EDTA or 1 mM CaCl_2_ (for apo- or 2Ca^2+^-conformer, respectively), or 10 mM Tris-HCl pH 7.4, 50 мМ KCl, 20 μМ DTT, 100 μM ZnCl_2_ or 2 μM CaCl_2_ or 2 μM CaCl_2_ and 100 μM ZnCl_2_ or 1 mM CaCl_2_ and 100 μM ZnCl_2_ (for Zn^2+^- or 1Ca^2+^- or 1Ca^2+^/Zn^2+^- or 2Ca^2+^/Zn^2+^-conformers, respectively). A contribution of the buffer was subtracted from experimental spectra. Band width was 2 nm, averaging time 2 s, and accumulation 3. Quantitative estimates of the secondary structure contents were carried out using SELCON3, CDSSTR and CONTIN algorithms, and SDP48 and SMP56 reference protein sets, as implemented in CDPro software package ^68^. The final secondary structure fractions represent averaged values.

### 5.4. Intrinsic (tryptophan) fluorescence spectroscopy (IFS)

Calcium binding to myristoylated forms of hRec (1.7 μM) and bRec (1.9 μM) at 25 °C was monitored by changes in its intrinsic fluorescence, excited at 280 nm, using a Cary Eclipse spectrofluorometer (Varian Inc., Palo Alto, CA, USA) equipped with a cuvette holder controlled by a Peltier element. Measurements were performed in quartz cuvettes with a path length of 10 mm in 20 mM MES-HCl pH 6.6-6.8, 50 mM KCl, 20 μM DTT, containing 1.9-2.0 mM DTPA-KOH and various aliquots of CaCl_2_ standard solution to achieve the required free calcium concentration, calculated using the IBFluo v.1.1 program (IBI RAS, Pushchino, Russia). The dependence of protein fluorescence intensity on free Ca^2+^ concentration was approximated by a Hill model ^69^.

### 5.5. Equilibrium Dialysis/Atomic Absorption Spectroscopy (AAS)

The binding of zinc to myristoylated forms of hRec at (25.5±0.5) °C was studied using atomic absorption spectroscopy coupled to equilibrium dialysis according to a previously developed protocol ^64^. The dialysis was performed in a teflon block (HT-Dialysis, Gales Ferry, CT, USA) with a regenerated cellulose membrane (3.5 kDa MWCO). 50 µL of 10-12 µM solutions of hRec and (0.5 to 122) µM of ZnCl_2_ in 25 mM Hepes-KOH, 50 mM KCl, pH 7.4 were placed on both sides of the dialysis membrane and equilibrated under continuous mixing at 140 rpm for 6 h. The concentration of Zn^2+^ in each half of the wells was determined using an atomic absorption spectrometer Thermo Scientific iCE 3000 with electrothermal atomization equipped with Zeeman background correction (Thermo Scientific, Hemel Hempstead, UK) at the absorption band of 213.9 nm. The analytical signal was calibrated using AAS standard solution of Zn^2+^ (Sigma-Aldrich, St. Louis, MO, USA). The concentration of free Zn^2+^ was assumed to be equal to that measured in the half of the well without recoverin, and the concentration of Zn^2+^ bound to the protein was estimated as the difference between the total metal concentrations in both halves of the well.

### 5.6. Equilibrium Centrifugation Assay

The membrane-binding properties of myristoylated forms of hRec and bRec were studied using ROS membranes obtained from frozen bovine retinas and washed from peripheral proteins with urea ^22,23^ by equilibrium centrifugation assay, as described in ^14,64^ with modifications. Briefly, 5 μL of ROS suspension were mixed with 42 μM recoverin in 20 mM Tris-HCl buffer (pH 8.0), 150 mM NaCl, 1 mM DTT, 20 mM MgCl_2_, in the presence of Ca^2+^-buffers prepared from 1 mM diBr-BAPTA and 0–1.5 mM CaCl_2_ (yielding 0.1-500 μM [Ca^2+^]_free_). Alternatively, Ca^2+^-buffers were replaced by one of the following components: 1 mM EDTA (apo form), 10 µM CaCl_2_ (1Ca^2+^ bound form), 1 mM CaCl_2_ (2Ca^2+^ bound form), 10 µM CaCl_2_/50 µM ZnCl_2_ (1Ca^2+^/Zn^2+^ bound form), or 1 mM CaCl_2_/50 µM ZnCl_2_ (2Ca^2+^/Zn^2+^ bound form). After incubation in a thermostatic shaker for 30 minutes (37 °C, 1000 rpm), the membrane pellet was collected by centrifugation (24,000 g, 15 minutes) and dissolved in 50 μl of SDS-PAGE sample buffer. Membrane-bound bRec or hRec was visualized by SDS-PAGE with Coomassie Brilliant Blue staining, and their weight fractions were quantified by densitometric analysis of protein bands using GelAnalyzer software (www.gelanalyzer.com).

### 5.7. Surface Plasmon Resonance (SPR)

The interaction between myristoylated forms of hRec/bRec and GST-GRK1-25 was analyzed using surface plasmon resonance (SPR) biosensor technology on a Bio-Rad ProteOn XPR36 instrument (Bio-Rad, Hercules, CA, USA). The GST-GRK1-25 and GST (control) ligands were immobilized on the ProteOn GLH chip by amine coupling, reaching SPR signals of 10,000 resonance units (RUs) and 10,500 RUs, respectively. Various concentrations of hRec or bRec (analytes) in 10 mM HEPES, 150 mM NaCl, 0.05% Tween 20, pH 7.4 (running buffer) containing either 1 mM EDTA or 1 mM CaCl_2_ and/or 30/60/90 µM ZnCl_2_ (for hRec/bRec concentrations 0-10/20/30 µM, respectively) were applied to the chip at a flow rate of 30 µL/min. The association phase was 300 s. Complex formation was monitored in a series of titrations in which the hRec/bRec sample dialyzed against the running buffer was used at concentrations from 0.6/0.4 μM to 40/30 μM, respectively, in the presence or absence of Ca^2+^ and/or Zn^2+^ in the buffer. The signal of GST was subtracted from the sensorgrams, and the maximal amplitudes of the response were plotted as a function of recoverin concentration. The equilibrium analysis of SPR data was performed within the Langmuir model using ProteOn Manager software (Bio-Rad).

### 5.8. Crystallization

Non-myristoylated hRec (476 µM) was crystallized using sitting drop vapor diffusion at 20°C. The stock protein solutions in 40 mM Tris-HCl, pH 8.0, 150 mM NaCl, 1 mM DTT, 5 mM ZnCl₂, 2 mM TCEP were screened using the commercial SG1 screens (Molecular Dimensions, USA) with an NT8 robotic system (Formulatrix, USA). Crystals (Fig. S4) were obtained within a week under the following conditions: 82% saturated sodium citrate, 5 mM CaCl₂, pH 7.0 for the 2Ca^2+^-bound form and 2.0 M ammonium sulfate, 0.1 M Bis-tris, pH 5.5 for the 1Ca^2+^/2Zn^2+^-bound form.

### 5.9. X-ray Data Collection and Structure Determination

The crystals of non-myristoylated hRec were flash-cooled in liquid nitrogen prior to data collection. X-ray diffraction data were collected at 100 K at the Shanghai Synchrotron Radiation Facility (SSRF, Shanghai, China) using a wavelength of 0.979 Å. The 2Ca²⁺-bound and 1Ca^2+^/2Zn^2+^-bound datasets were obtained at beamlines BL10U2 and BL17U1, respectively. Diffraction data were processed and scaled using XDS software package tool ^70^. The 1Ca^2+^/2Zn^2+^ dataset exhibited strong anomalous scattering (Table 2). Molecular replacement was performed using a polyalanine model derived from truncated mutant human recoverin (PDB ID: 2D8N) as the search model. Iterative model building and refinement were conducted in the Phenix ^71^ Suite program and COOT ^72^. The 2Ca²-bound structure was refined with anisotropic atomic displacement parameters, and the 1Ca^2+^/2Zn^2+^-bound structure employed TLS refinement. Both structures crystallized in space group I4, with one molecule per asymmetric unit. Calcium ions were identified in electron density maps, while anomalous difference maps unambiguously enabled positioning two zinc ions in the 1Ca^2+^/2Zn^2+^ model. Metal coordination geometries were validated using the CheckMyMetal web server ^73^. Model quality was assessed with phenix.molprobity ^74^ and the Quality Control Check web server (https://qc-check.usc.edu). Data collection and refinement statistics are detailed in Table 2. The structural figures were generated using PyMOL (version 2.5.0).

### 5.10. Molecular Dynamics (MD) Simulations and Free Energy Calculation

The initial 2Ca²-bound structure was prepared by modeling missing side chains and reconstructing the distorted N-terminal region. A myristoyl group was added to the protein, which was subsequently embedded in a phospholipid bilayer containing 262 lipids (DOPC: DOPG, 4:1 ratio ^12^) using the CHARMM-GUI server ^75^. The system was solvated with approximately 21,000 TIP3P water molecules and neutralized with 150 mM KCl, resulting in a simulation box of 101,264 atoms. Initial lipid penetration was assessed to prevent artifactual protein-membrane contacts. Minimization and equilibration steps followed the standard CHARMM-GUI protocol with gradually released positional restraints on protein and membrane heavy atoms. Production simulation was run for 500 ns without restraints. All simulations were carried out using GROMACS 2023.1 ^76^. Temperature (310 K) and pressure (1 bar) were controlled using the V-rescale thermostat and C-rescale semi-isotropic barostat with coupling time constants of 0.5 ps and 5.0 ps. The integration time step was 1 fs during the initial equilibration and 2 fs for subsequent equilibration and production. Bonds to hydrogens were constrained using LINCS ^77^. Long-range electrostatics were treated with PME, and short-range interactions with the Verlet cutoff scheme. The CHARMM36m force field ^78^ was used for proteins and lipids, and the TIP3P model for water.

SASA was computed from the production trajectory using the Shrake–Rupley algorithm implemented in MDTraj ^79^. Residues showing a significant SASA difference in the presence and in the absence of membrane were considered membrane-contacting.

Hybrid topologies for each mutation were generated using pmx ^80^. For alchemical transformations that changed the net charge of the system, an appropriate counterion was included in the hybrid topology and its charge was gradually turned off during the transformation ^81^. Thermodynamic integration simulations were performed for each mutation in both the membrane-bound and membrane-free systems. The potential energy function interpolated between the end states A and B as: H(λ)=HA−(1−λ)HB. Both VdW and electrostatic terms were coupled/decoupled simultaneously. The following soft-core parameters were used: sc-alpha = 0.5, sc-power = 1, sc-sigma = 0.3. λ-windows were simulated at λ = 0, 0.02, 0.05, 0.08, 0.12, 0.16, 0.2, 0.25, 0.3, 0.4, 0.5, 0.6, 0.7, 0.78, 0.85, 0.9, 0.94, 0.97, 0.99, and 1.0, with 5 ns per window. The free energy ΔG was computed by numerical integration of the ⟨dH/dλ⟩ values using the trapezoidal rule (NumPy implementation). ΔΔG values were obtained through a thermodynamic cycle comparing membrane-bound and membrane-free transformations. The uncertainty of the calculated free energy change was estimated using a block-averaging approach applied to the ⟨∂H/∂λ⟩ values from each λ-window. For every simulation, the derivative time series was divided into equal temporal blocks, and the mean derivative within each block was computed. The standard error of the mean (SEM) was then obtained as the standard deviation of these block means divided by the square root of the number of blocks, providing an estimate of statistical uncertainty that accounts for temporal correlations in the trajectory. The individual SEM values from all λ-windows were propagated to yield the overall error of the integrated free energy difference.

### 5.11. X-ray Absorption Near Edge Structure (XANES) analysis

#### 5.11.1. Experimental Procedure

The experiments were conducted at the LANGMUIR beamline, Kurchatov Center for Synchrotron Radiation (Moscow, Russia) using protein monolayers as described in ^82^. The monolayers were prepared from 0.1 μM hRec or bRec, preincubated with 50 μM CaCl_2_ and 50 μM ZnCl_2_, and formed in a 35 ml Teflon trough on the surface of a buffered liquid subphase (50 mM Tris-HCl, pH 7.5). To qualitatively confirm the ability of Ca^2+^-loaded recoverin orthologues to bind zinc, characteristic X-ray fluorescence spectra (Fig. 1E) were recorded at the fixed angle 0.8×θ_C_, where θ_C_ denotes the critical angle of total external reflection (TER) on water. The fluorescence was detected by an energy-dispersive Vortex EX detector positioned at 90° above the liquid surface. XANES spectra (Fig. 3B) at the Zn K-edge were collected in fluorescence mode under TER conditions. Energy calibration was performed with reference to the Zn K-edge of a metallic zinc foil. To improve high-statistic XANES spectra, several energy scans were recorded and averaged using multipass data acquisition. The reproducibility of the energy position of the monochromator was determined to be within 0.18 eV. During the measurements, the enclosed Teflon trough was purged continuously with helium saturated with water vapor to minimize X-ray scattering and reduce evaporation from the liquid subphase. All measurements were performed at 25°C.

#### 5.11.2. Calculations

##### 5.11.2.1. Geometry Optimization using Quantum Mechanics/Molecular Mechanics (QM/MM) approach

To optimize the coordination geometry of the zinc ions within the protein, we performed QM/MM calculations on the zinc-bound structure. Protonation states of titratable residues were first assigned using the PropKa web server ^83,84^, and hydrogens were added with the pdb4amber tool from the AmberTools package ^85^. Potential steric clashes were removed through energy minimization using the ff14SB AMBER force field ^86^. Geometry optimizations were performed with an electrostatic embedding scheme QM/MM scheme ^87^ implemented in Orca v6.1.0 ^88^, using the L-BFGS ^89^ algorithm for efficient convergence. The QM region comprised the side chains of all residues within 6 Å of each zinc ion, while the MM region included all atoms within 6 Å of the QM region. Remaining protein atoms were held fixed during optimization to maintain global structural integrity. The QM subsystem was treated at the density functional theory level using B3LYP ^90^ functional, augmented by Grimme’s D3 dispersion correction ^91^ with Becke-Johnson damping ^92^. The correlation-consistent cc-pVDZ basis set ^93^ was applied. To accelerate the self-consistent field procedure, the Resolution–of–Identity ^94^ approximation was used, with auxiliary basis sets matching the atomic basis. The MM region was treated using FF14SB AMBER force field ^95^.

##### 5.11.2.2. Spectra Calculations

Theoretical XAS spectra on the K-edge of zinc were calculated using the FDMNES code ^96,97^. The exchange-correlation potential was parameterized following the Hedin and Lundqvist formalism^98^. Unoccupied electronic states were determined by solving the Schrödinger equation employing the finite difference method (FDM). Although FDM is more computationally demanding than the muffin-tin approximation, it has demonstrated superior accuracy for metal-organic systems, as evidenced in a previous study ^99^. To ensure close agreement with experimental data, convolution parameters for the theoretical spectra were iteratively optimized to minimize deviations from the measured XANES spectra. All calculations were performed using the QM/MM-optimized structural models, with clusters defined by a 6 Å radius around the zinc coordination site to capture the local electronic environment accurately (Fig. 3B).

### 5.12. Data Processing and Statistics

The experimental data were visualized and analyzed using SigmaPlot 11 (Systat Software, Erkrath, Germany) and Microcal OriginPro 9.0 (OriginLab Corporation, Northampton, MO, USA) software. The graphs and histograms in the figures represent the mean ± SEM. Each experiment was performed in at least 3 replicates.

## Data Availability

The hRec structural coordinates for 1Ca^2+^/2Zn^2+^ and 2Ca^2+^-bound conformers are deposited in RCSB as 24JC and 24JB, respectively. All scripts, input files, and MD analysis workflows are available in Zenodo repository 10.5281/zenodo.19050052.

## Acknowledgments

N.N.N., V.I.B. and E.Y.Z. acknowledge the Ministry of Science and Higher Education of the Russian Federation (agreement # 075-15-2025-512; determination of the crystal structure of Ca^2+^-saturated hRec and XANES analysis on the LANGMUIR beamline at the Kurchatov Synchrotron Radiation Center (Moscow, Russia)). We thank SSRF for providing the opportunity to collect crystallographic data at the beamlines BL10U2 and BL17UM. The study of zinc affinity and zinc-dependent structural properties of apo- and Ca^2+^-loaded conformers of hRec, as well as X-ray diffraction analysis of the 1Ca^2+^-bound hRec complex with zinc (1Ca^2+^/Zn^2+^-bound form), were supported by the Russian Science Foundation grant No. 24-15-00171 (to E.Y.Z.). D.A.F. acknowledges the Russian Science Foundation grants No. 25-74-00077 and 23-14-00160 for support of computational time. D.A.F. acknowledges the use of the High Performance Computing Facility at Lobachevsky State University and SKIF (Siberian Circular Photon Source), and associated support services. P.S.O. is a member of Guangdong Provincial Higher Education Institution Innovation Team Project (Grant #2025KCXTD055). The development and validation of the procedure for analyzing conformational stability using nanoDSF and its adaptation for studying NCS proteins were supported by the Russian Science Foundation (A.V.M., D.E.D., project No. 22-74-10036-П). M.L.S., N.G.S., and E.Y.Z. are grateful to the Moscow State University Development Program for providing access to the nanoDSF instrument PSA-16. A.S.B. acknowledges the support of the Ministry of Science and Higher Education under agreement No. 075-03-2026-305 (project FSMG-2024-0012) for studies of membrane protein interactions with the lipid bilayer. The analysis of general principles of functioning of photoreceptor proteins, including recoverin, is carried out under the state assignment of Lomonosov Moscow State University (M.L.S., N.G.S., and E.Y.Z.).

## Authors Contributions

Project Conceptualization – S.E.P.; V.I.B., E.Y.Z.; Methodology – S.E.P.; V.I.B., E.Y.Z.; Investigation – C.O.M., A.A.V., A.S.B., N.N.N., V.A.R., M.P.S., M.L.S., N.G.S., M.B.S., I.A.K., A.V.M., D.E.D., Y.Y., D.A.F., E.A.L., E.I.P., D.V.Z., A.L.T., A.V.R., S.N.Y., P.S.O., V.I.B.; Visualization – C.O.M., P.S.O. and E.Y.Z.; Writing Original Manuscript – C.O.M. and E.Y.Z.; Writing – Review & Editing: S.E.P., V.I.B., E.Y.Z.; Funding Acquisition – N.N.N., V.I.B., E.Y.Z.; Project Supervision – S.E.P., V.I.B., E.Y.Z.

