## Supplementary File for "Structural insights into human recoverin"

### Slide 1
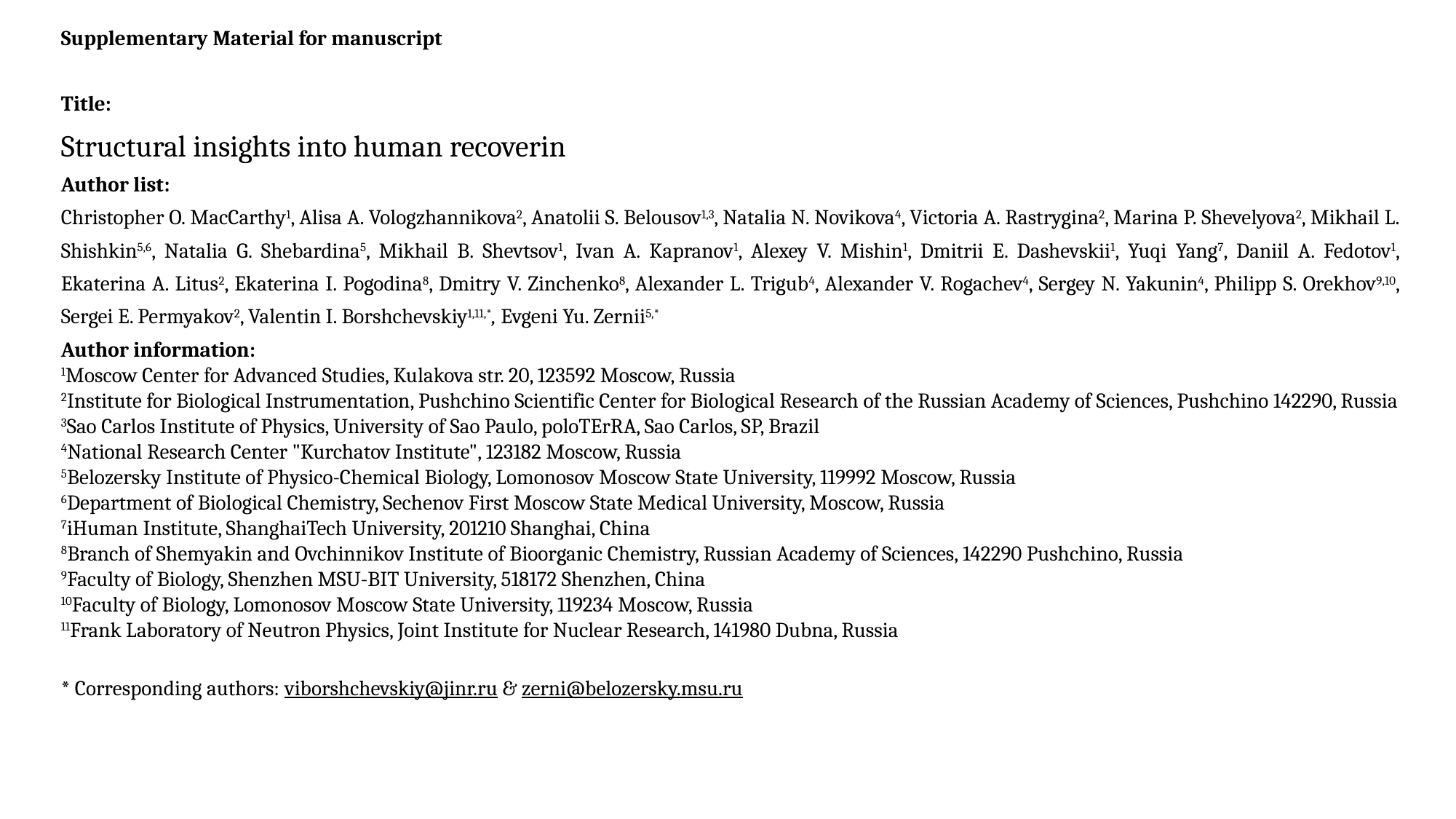

Supplementary Material for manuscript
Title:
Structural insights into human recoverin
Author list:
Christopher O. MacCarthy^1^, Alisa A. Vologzhannikova^2^, Anatolii S. Belousov^1,3^, Natalia N. Novikova^4^, Victoria A. Rastrygina^2^, Marina P. Shevelyova^2^, Mikhail L. Shishkin^5,6^, Natalia G. Shebardina^5^, Mikhail B. Shevtsov^1^, Ivan A. Kapranov^1^, Alexey V. Mishin^1^, Dmitrii E. Dashevskii^1^, Yuqi Yang^7^, Daniil A. Fedotov^1^, Ekaterina A. Litus^2^, Ekaterina I. Pogodina^8^, Dmitry V. Zinchenko^8^, Alexander L. Trigub^4^, Alexander V. Rogachev^4^, Sergey N. Yakunin^4^, Philipp S. Orekhov^9,10^, Sergei E. Permyakov^2^, Valentin I. Borshchevskiy^1,11,*^, Evgeni Yu. Zernii^5,*^
Author information:
^1^Moscow Center for Advanced Studies, Kulakova str. 20, 123592 Moscow, Russia
^2^Institute for Biological Instrumentation, Pushchino Scientific Center for Biological Research of the Russian Academy of Sciences, Pushchino 142290, Russia
^3^Sao Carlos Institute of Physics, University of Sao Paulo, poloTErRA, Sao Carlos, SP, Brazil
^4^National Research Center "Kurchatov Institute", 123182 Moscow, Russia
^5^Belozersky Institute of Physico-Chemical Biology, Lomonosov Moscow State University, 119992 Moscow, Russia
^6^Department of Biological Chemistry, Sechenov First Moscow State Medical University, Moscow, Russia
^7^iHuman Institute, ShanghaiTech University, 201210 Shanghai, China
^8^Branch of Shemyakin and Ovchinnikov Institute of Bioorganic Chemistry, Russian Academy of Sciences, 142290 Pushchino, Russia
^9^Faculty of Biology, Shenzhen MSU-BIT University, 518172 Shenzhen, China
^10^Faculty of Biology, Lomonosov Moscow State University, 119234 Moscow, Russia
^11^Frank Laboratory of Neutron Physics, Joint Institute for Nuclear Research, 141980 Dubna, Russia

### Slide 2
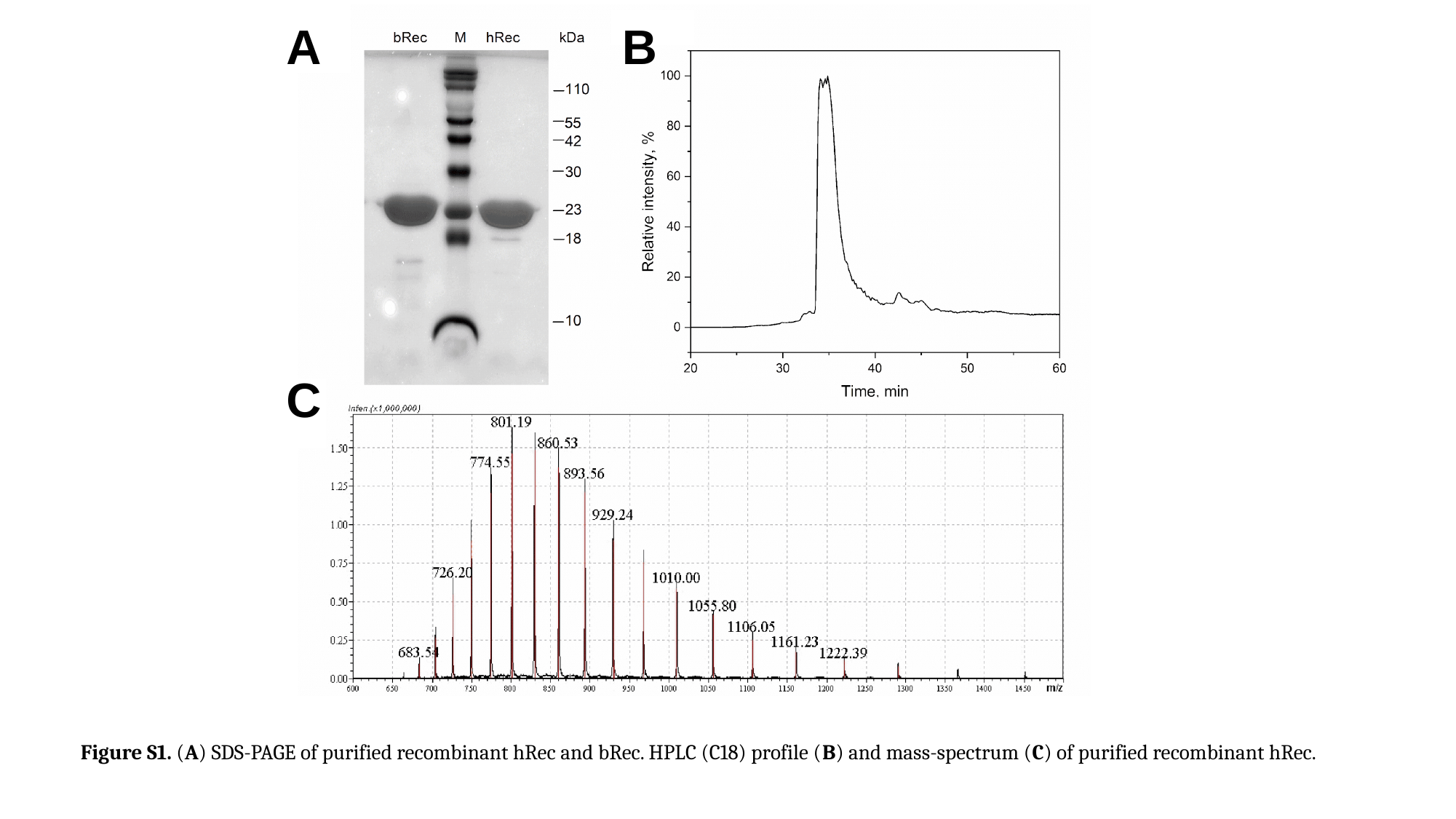

A
B
C
Figure S1. (A) SDS-PAGE of purified recombinant hRec and bRec. HPLC (C18) profile (B) and mass-spectrum (C) of purified recombinant hRec.

### Slide 3
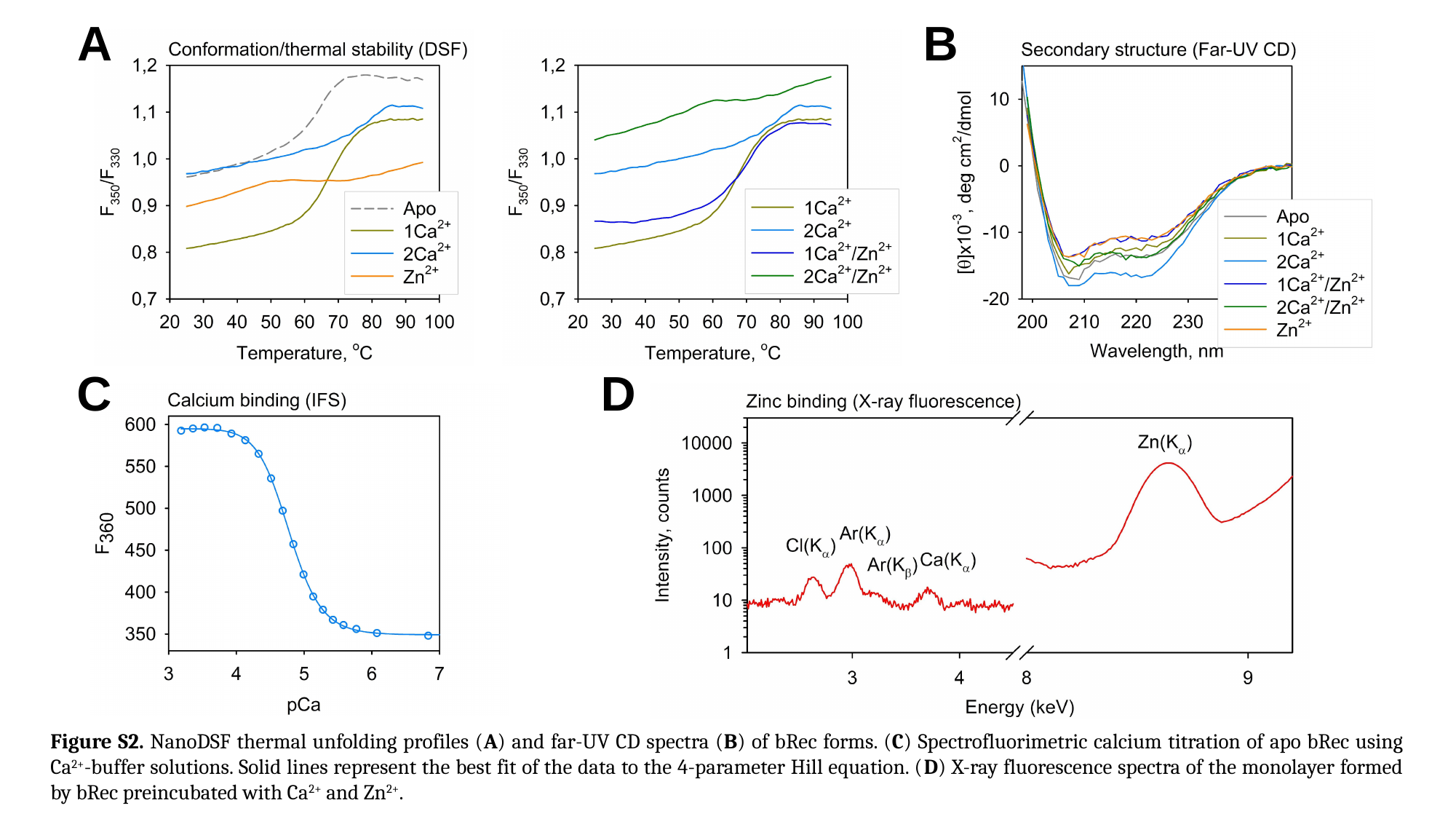

A
B
С
D
Figure S2. NanoDSF thermal unfolding profiles (A) and far-UV CD spectra (B) of bRec forms. (C) Spectrofluorimetric calcium titration of apo bRec using Ca^2+^-buffer solutions. Solid lines represent the best fit of the data to the 4-parameter Hill equation. (D) X-ray fluorescence spectra of the monolayer formed by bRec preincubated with Ca^2+^ and Zn^2+^.

### Slide 4
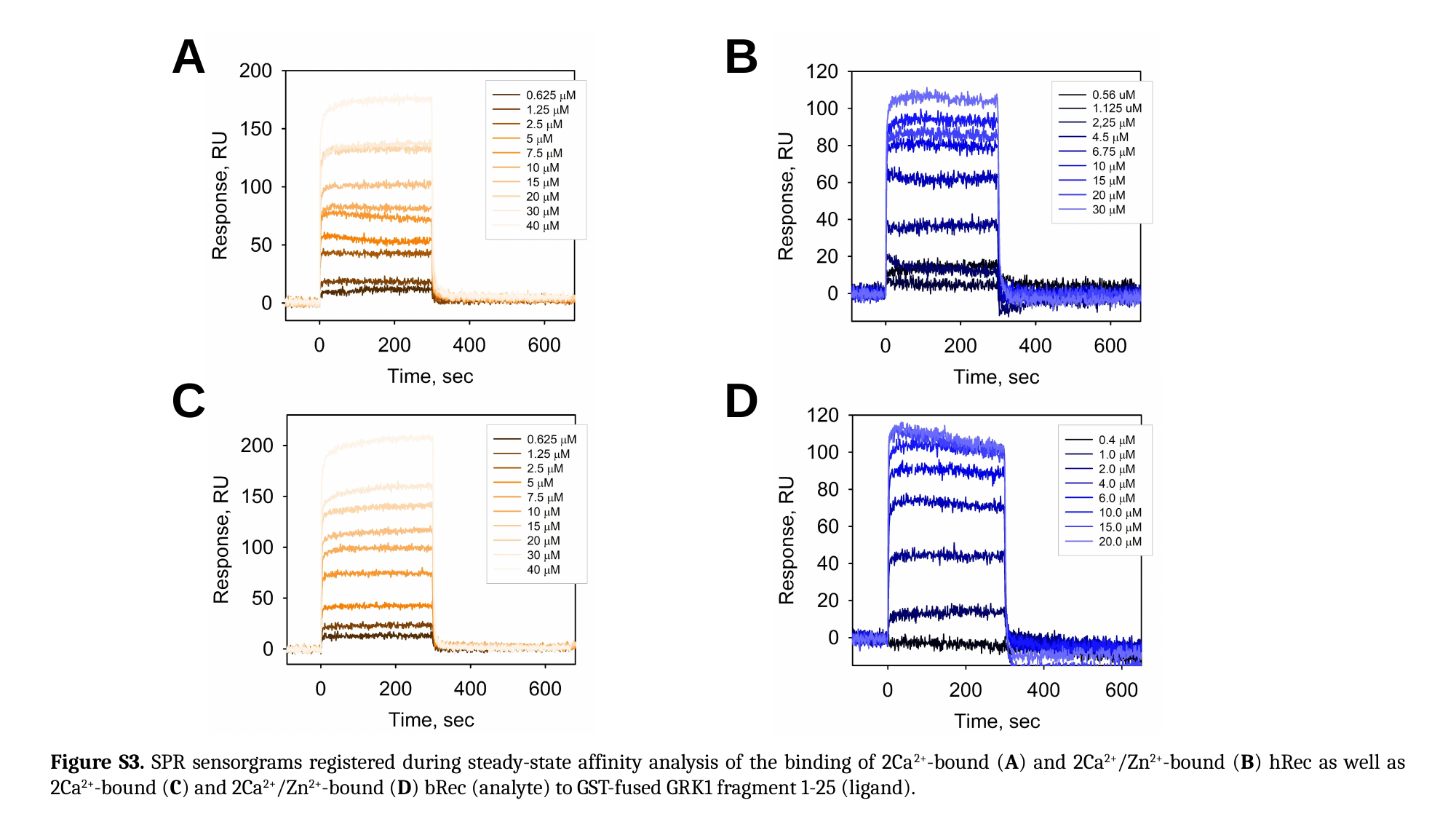

A
B
C
D
Figure S3. SPR sensorgrams registered during steady-state affinity analysis of the binding of 2Ca^2+^-bound (A) and 2Ca^2+^/Zn^2+^-bound (B) hRec as well as 2Ca^2+^-bound (C) and 2Ca^2+^/Zn^2+^-bound (D) bRec (analyte) to GST-fused GRK1 fragment 1-25 (ligand).

### Slide 5
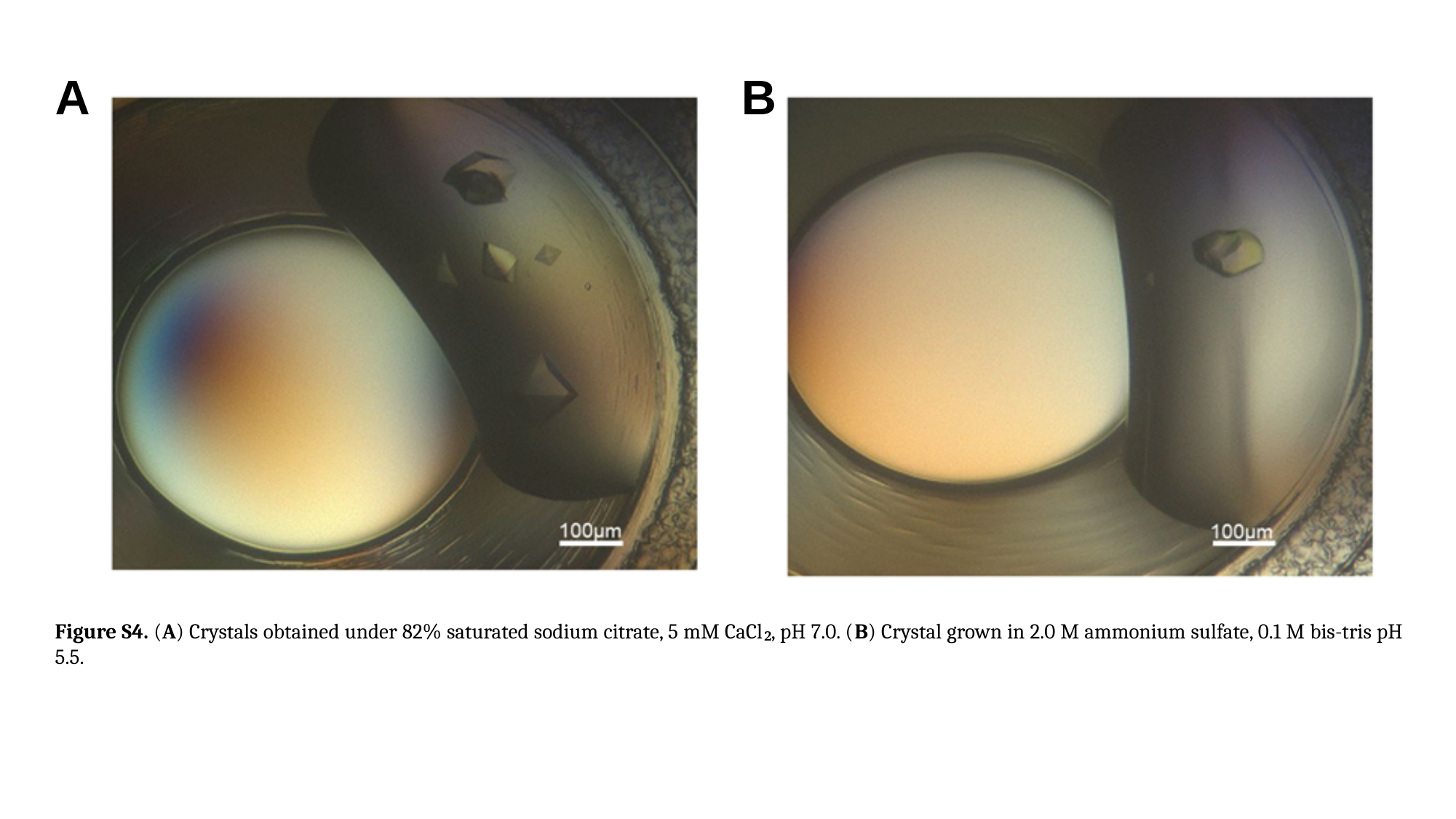

A
B
Figure S4. (A) Crystals obtained under 82% saturated sodium citrate, 5 mM CaCl₂, pH 7.0. (B) Crystal grown in 2.0 M ammonium sulfate, 0.1 M bis-tris pH 5.5.

### Slide 6
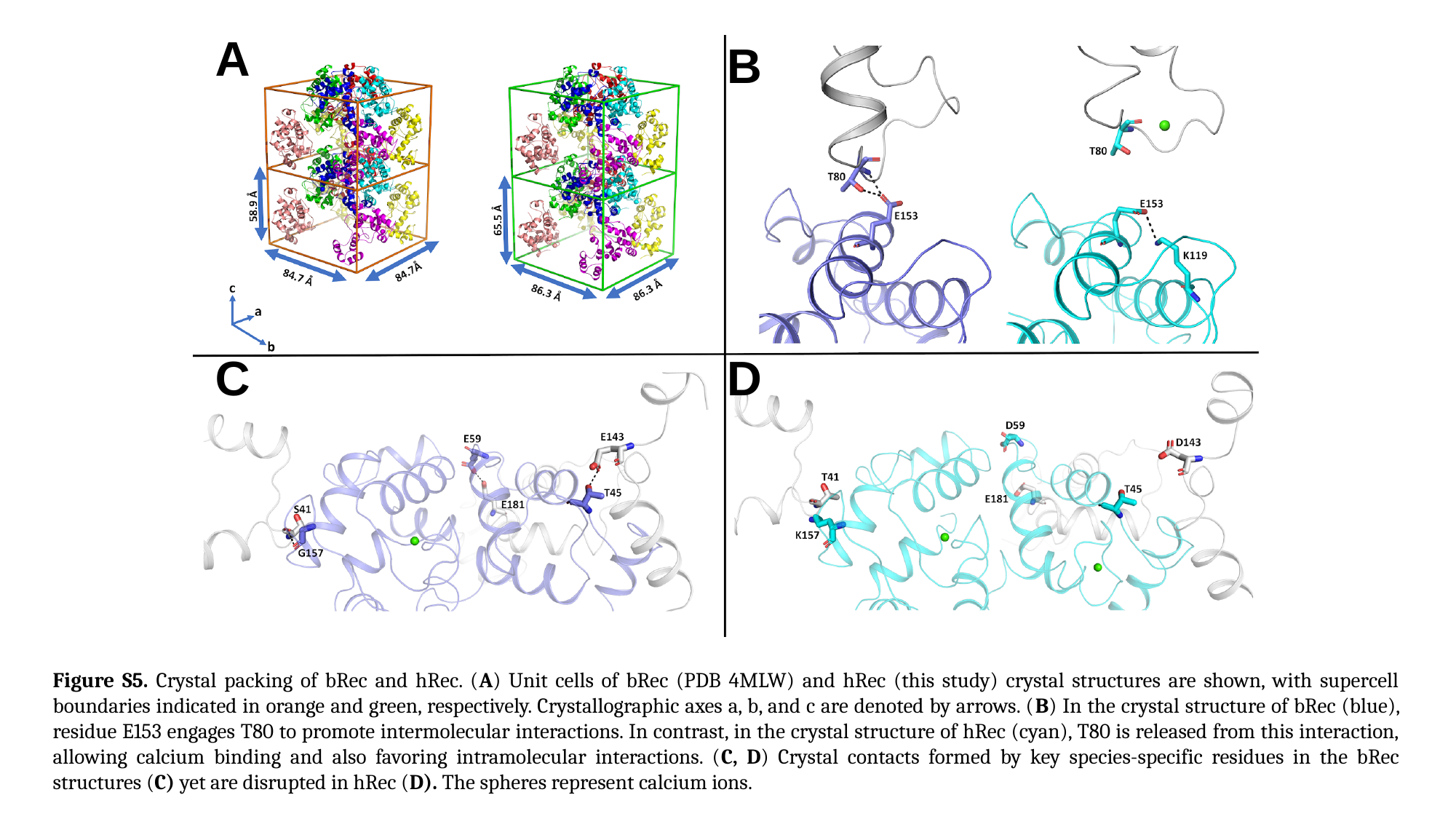

A
B
С
D
Figure S5. Crystal packing of bRec and hRec. (A) Unit cells of bRec (PDB 4MLW) and hRec (this study) crystal structures are shown, with supercell boundaries indicated in orange and green, respectively. Crystallographic axes a, b, and c are denoted by arrows. (B) In the crystal structure of bRec (blue), residue E153 engages T80 to promote intermolecular interactions. In contrast, in the crystal structure of hRec (cyan), T80 is released from this interaction, allowing calcium binding and also favoring intramolecular interactions. (C, D) Crystal contacts formed by key species-specific residues in the bRec structures (C) yet are disrupted in hRec (D). The spheres represent calcium ions.

### Slide 7
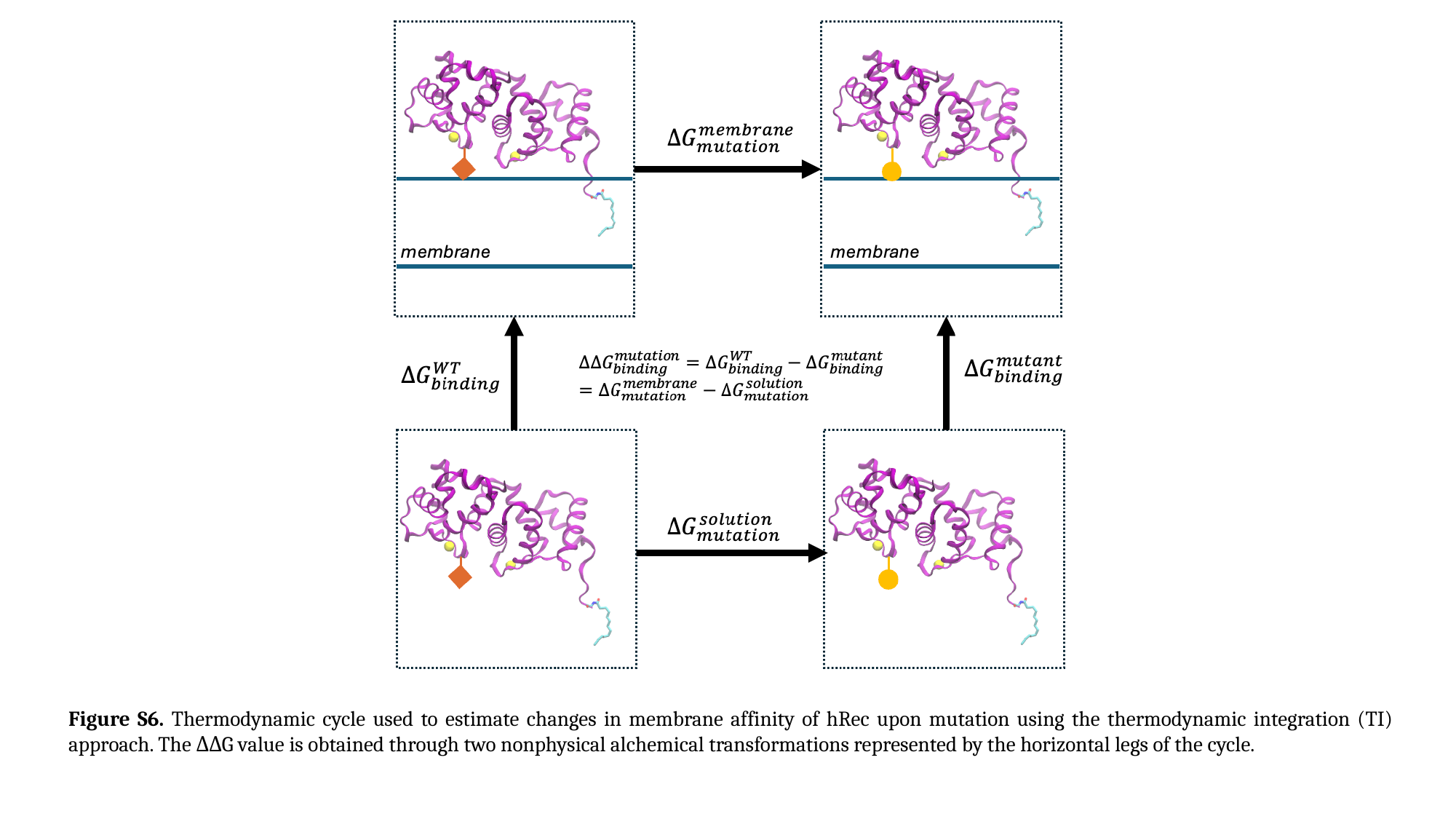

Figure S6. Thermodynamic cycle used to estimate changes in membrane affinity of hRec upon mutation using the thermodynamic integration (TI) approach. The ΔΔG value is obtained through two nonphysical alchemical transformations represented by the horizontal legs of the cycle.

### Slide 8
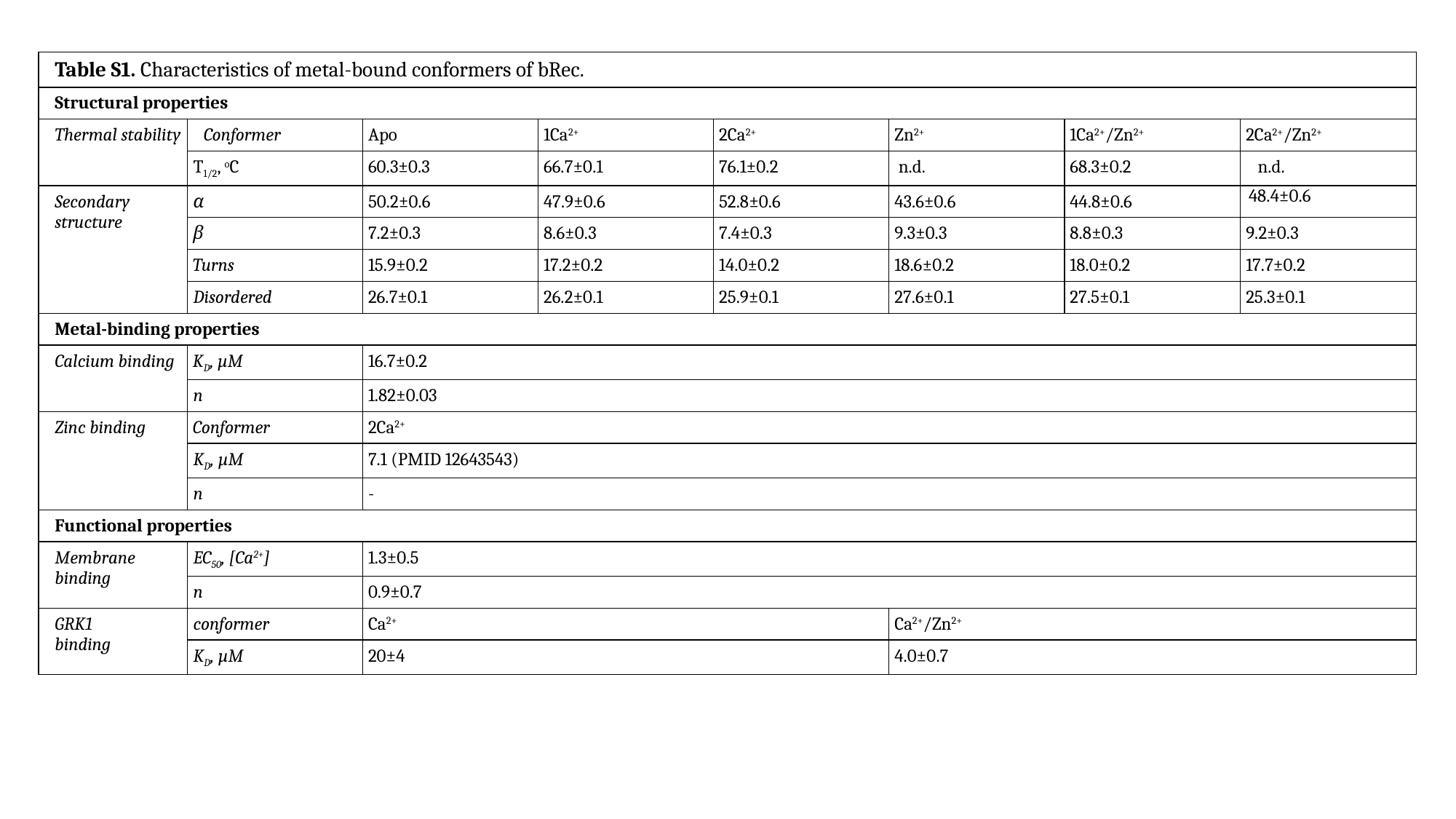

| Table S1. Characteristics of metal-bound conformers of bRec. | | | | | | | |
| --- | --- | --- | --- | --- | --- | --- | --- |
| Structural properties | | | | | | | |
| Thermal stability | Conformer | Apo | 1Ca^2+^ | 2Ca^2+^ | Zn^2+^ | 1Ca^2+^/Zn^2+^ | 2Ca^2+^/Zn^2+^ |
| | T_1/2_, ^o^C | 60.3±0.3 | 66.7±0.1 | 76.1±0.2 | n.d. | 68.3±0.2 | n.d. |
| Secondary structure | α | 50.2±0.6 | 47.9±0.6 | 52.8±0.6 | 43.6±0.6 | 44.8±0.6 | 48.4±0.6 |
| | β | 7.2±0.3 | 8.6±0.3 | 7.4±0.3 | 9.3±0.3 | 8.8±0.3 | 9.2±0.3 |
| | Turns | 15.9±0.2 | 17.2±0.2 | 14.0±0.2 | 18.6±0.2 | 18.0±0.2 | 17.7±0.2 |
| | Disordered | 26.7±0.1 | 26.2±0.1 | 25.9±0.1 | 27.6±0.1 | 27.5±0.1 | 25.3±0.1 |
| Metal-binding properties | | | | | | | |
| Calcium binding | K_D_, µM | 16.7±0.2 | | | | | |
| | n | 1.82±0.03 | | | | | |
| Zinc binding | Conformer | 2Ca^2+^ | | | | | |
| | K_D_, µM | 7.1 (PMID 12643543) | | | | | |
| | n | - | | | | | |
| Functional properties | | | | | | | |
| Membrane binding | EC_50_, [Ca^2+^] | 1.3±0.5 | | | | | |
| | n | 0.9±0.7 | | | | | |
| GRK1 binding | conformer | Ca^2+^ | | | Ca^2+^/Zn^2+^ | | |
| | K_D_, µM | 20±4 | | | 4.0±0.7 | | |

### Slide 9
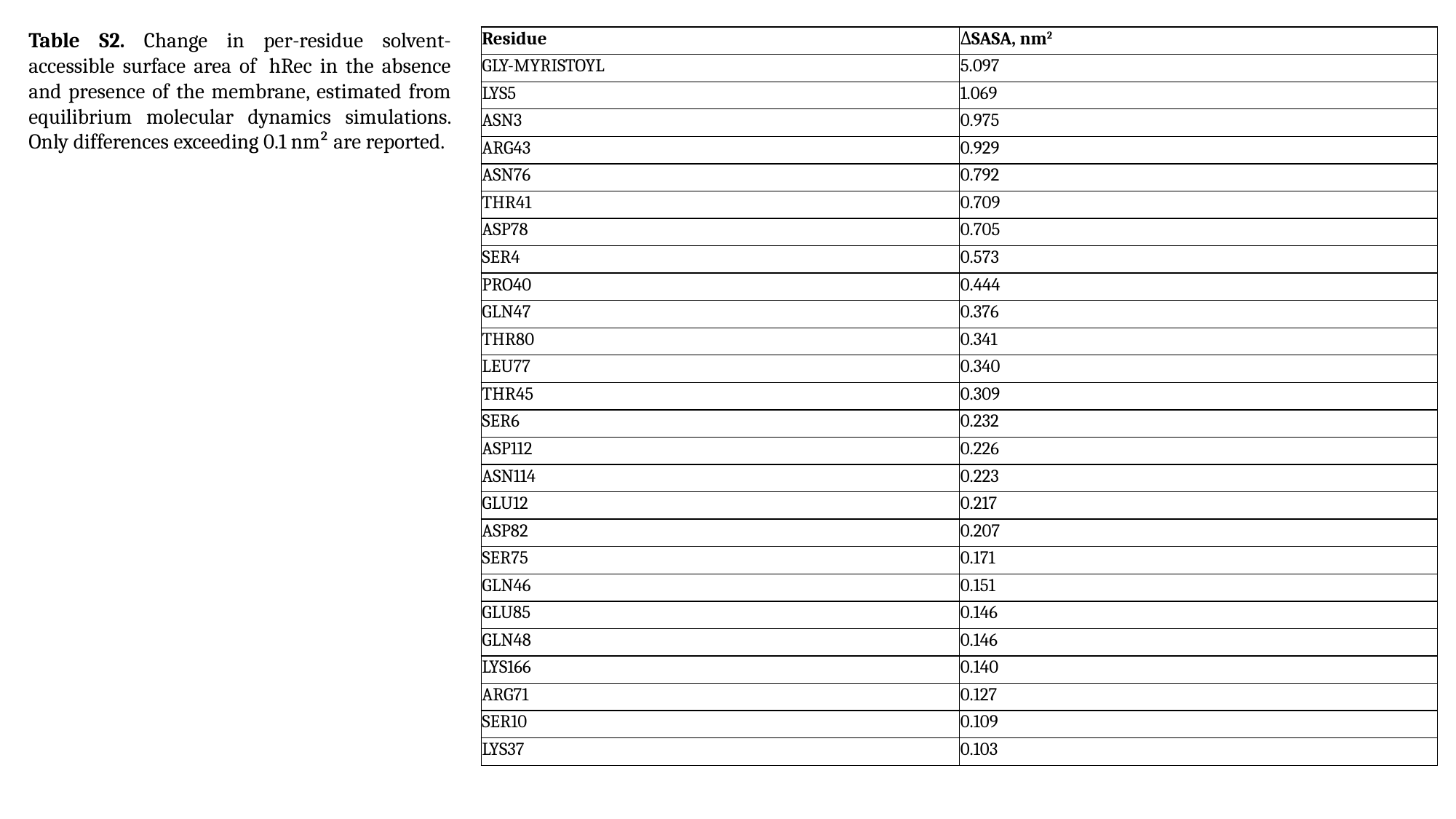

Table S2. Change in per-residue solvent-accessible surface area of  hRec in the absence and presence of the membrane, estimated from equilibrium molecular dynamics simulations. Only differences exceeding 0.1 nm² are reported.
| Residue | ΔSASA, nm^2^ |
| --- | --- |
| GLY-MYRISTOYL | 5.097 |
| LYS5 | 1.069 |
| ASN3 | 0.975 |
| ARG43 | 0.929 |
| ASN76 | 0.792 |
| THR41 | 0.709 |
| ASP78 | 0.705 |
| SER4 | 0.573 |
| PRO40 | 0.444 |
| GLN47 | 0.376 |
| THR80 | 0.341 |
| LEU77 | 0.340 |
| THR45 | 0.309 |
| SER6 | 0.232 |
| ASP112 | 0.226 |
| ASN114 | 0.223 |
| GLU12 | 0.217 |
| ASP82 | 0.207 |
| SER75 | 0.171 |
| GLN46 | 0.151 |
| GLU85 | 0.146 |
| GLN48 | 0.146 |
| LYS166 | 0.140 |
| ARG71 | 0.127 |
| SER10 | 0.109 |
| LYS37 | 0.103 |

### Slide 10
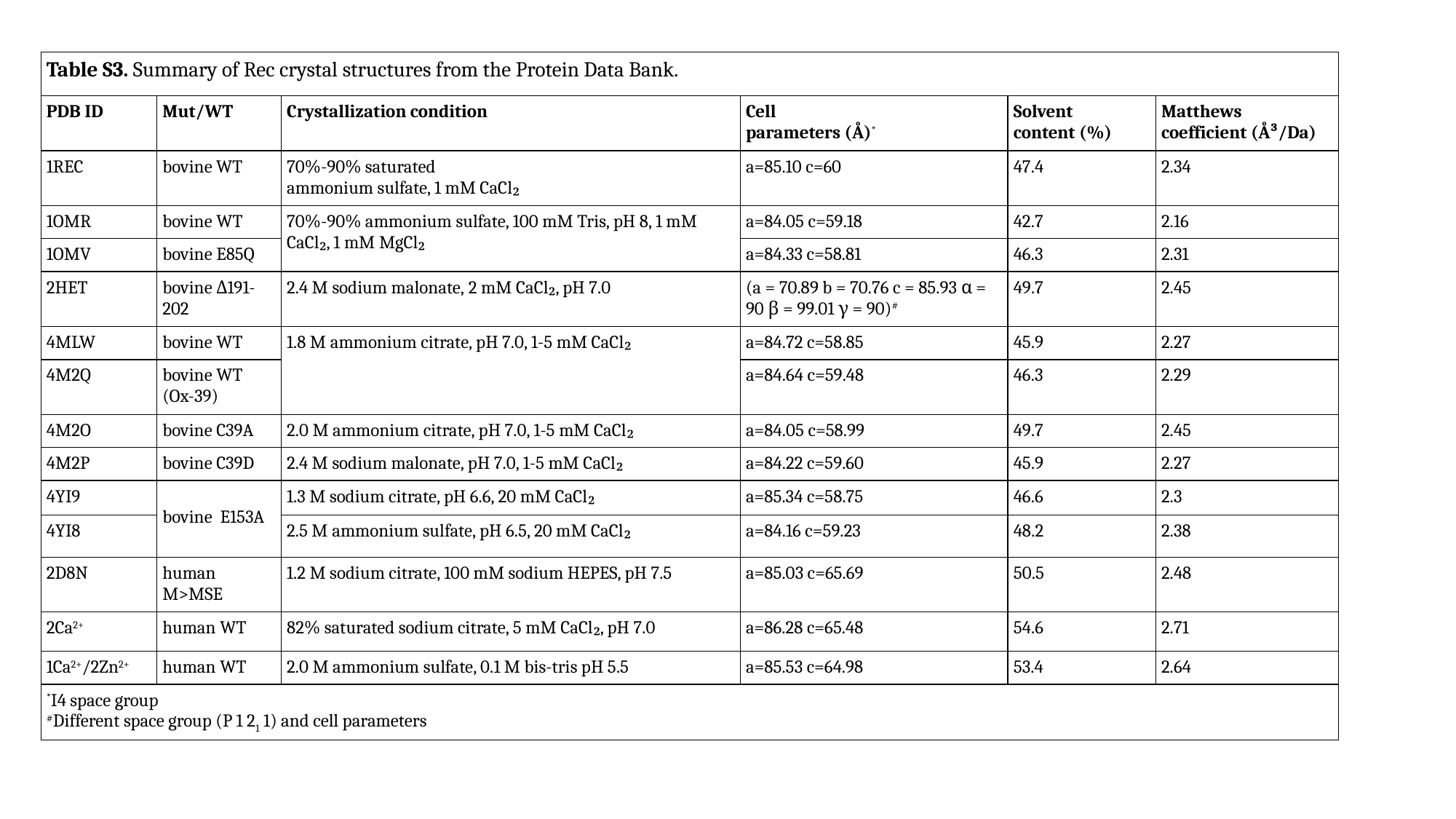

| Table S3. Summary of Rec crystal structures from the Protein Data Bank. | | | | | |
| --- | --- | --- | --- | --- | --- |
| PDB ID | Mut/WT | Crystallization condition | Cell parameters (Å)^\*^ | Solvent content (%) | Matthews coefficient (Å³/Da) |
| 1REC | bovine WT | 70%-90% saturated ammonium sulfate, 1 mM CaCl₂ | a=85.10 c=60 | 47.4 | 2.34 |
| 1OMR | bovine WT | 70%-90% ammonium sulfate, 100 mM Tris, pH 8, 1 mM CaCl₂, 1 mM MgCl₂ | a=84.05 c=59.18 | 42.7 | 2.16 |
| 1OMV | bovine E85Q | | a=84.33 c=58.81 | 46.3 | 2.31 |
| 2HET | bovine Δ191-202 | 2.4 M sodium malonate, 2 mM CaCl₂, pH 7.0 | (a = 70.89 b = 70.76 c = 85.93 α = 90 β = 99.01 γ = 90)^#^ | 49.7 | 2.45 |
| 4MLW | bovine WT | 1.8 M ammonium citrate, pH 7.0, 1-5 mM CaCl₂ | a=84.72 c=58.85 | 45.9 | 2.27 |
| 4M2Q | bovine WT (Ox-39) | | a=84.64 c=59.48 | 46.3 | 2.29 |
| 4M2O | bovine C39A | 2.0 M ammonium citrate, pH 7.0, 1-5 mM CaCl₂ | a=84.05 c=58.99 | 49.7 | 2.45 |
| 4M2P | bovine C39D | 2.4 M sodium malonate, pH 7.0, 1-5 mM CaCl₂ | a=84.22 c=59.60 | 45.9 | 2.27 |
| 4YI9 | bovine E153A | 1.3 M sodium citrate, pH 6.6, 20 mM CaCl₂ | a=85.34 c=58.75 | 46.6 | 2.3 |
| 4YI8 | | 2.5 M ammonium sulfate, pH 6.5, 20 mM CaCl₂ | a=84.16 c=59.23 | 48.2 | 2.38 |
| 2D8N | human M>MSE | 1.2 M sodium citrate, 100 mM sodium HEPES, pH 7.5 | a=85.03 c=65.69 | 50.5 | 2.48 |
| 2Ca^2+^ | human WT | 82% saturated sodium citrate, 5 mM CaCl₂, pH 7.0 | a=86.28 c=65.48 | 54.6 | 2.71 |
| 1Ca^2+^/2Zn^2+^ | human WT | 2.0 M ammonium sulfate, 0.1 M bis-tris pH 5.5 | a=85.53 c=64.98 | 53.4 | 2.64 |
| ^\*^I4 space group ^#^Different space group (P 1 2_1_ 1) and cell parameters | | | | | |
